# Elucidating Mechanisms of Antitumor Immunity Mediated by Live Oncolytic Vaccinia and Heat-Inactivated Vaccinia

**DOI:** 10.1101/2021.02.11.427912

**Authors:** Weiyi Wang, Peihong Dai, Shuaitong Liu, Ning Yang, Yi Wang, Rachel A. Giese, Taha Merghoub, Jedd D. Wolchok, Liang Deng

**Author notes:** corresponding author. Mailing address for Liang Deng: Dermatology Service, Department of Medicine, Memorial Sloan Kettering Cancer Center, 1275 York Ave., New York, NY 10065. These three authors contributed equally to this work (Weiyi Wang, Peihong Dai, Shuaitong Liu).

## Abstract

**Background:** Viral-based immunotherapy has the potential to overcome resistance to immune checkpoint blockade (ICB) and to fill the unmet needs of many cancer patients. Oncolytic viruses (OVs) are defined as engineered or naturally occurring viruses that selectively replicate in and kill cancer cells. OVs also induce antitumor immunity. The purpose of this study is to compare the antitumor effects of live OV-GM expressing murine granulocyte-macrophage colony-stimulating factor (mGM-CSF) versus inactivated OV-GM and elucidate the underlying immunological mechanisms.

**Methods:** In this study, we engineered a replication-competent, oncolytic vaccinia virus (OV-GM) by inserting a murine GM-CSF gene into the thymidine kinase (TK) locus of a mutant vaccinia E3LΔ83N, which lacks the Z-DNA-binding domain of vaccinia virulence factor E3. We compared the antitumor effects of intratumoral (IT) delivery of live OV-GM vs. heat-inactivated OV-GM (heat-iOV-GM) in a murine B16-F10 melanoma bilateral implantation model.

**Results:** Heat-iOV-GM infection of dendritic cells (DCs) and tumor cells in vitro induces type I IFN and pro inflammatory cytokines and chemokines, whereas live OV-GM does not. IT live OV-GM is less effective in generating systemic antitumor immunity compared with heat-iOV-GM. Similar to heat-iOV-GM, the antitumor effects of live OV-GM also require Batf3-dependent CD103+ dendritic cells. IT heat-iOV-GM induces higher numbers of infiltrating activated CD8+ and CD4+ T cells as well as higher levels of type I IFN, proinflammatory cytokines, and chemokines in the distant non-injected tumors, which is dependent on CD8+ T cells. When combined with systemic delivery of ICB, IT heat-iOV-GM is more effective in eradicating tumors compared with live OV-GM.

**Conclusions:** Tumor lysis induced by the replication of oncolytic DNA viruses has a limited effect on the generation of systemic antitumor immunity. The activation of Batf3-dependent CD103+ DCs is critical for antitumor effects induced by both live OV-GM and heat-iOV-GM. Heat-iOV-GM is more potent than live OV-GM in the induction of innate and adaptive immunity in both the injected and distant non-injected tumors. We propose that evaluations of both innate and adaptive immunity induced by IT oncolytic viral immunotherapy at injected and non-injected tumors should be included as potential biomarkers.

## Background

Oncolytic viral therapy has the potential to overcome resistance to immune checkpoint blockade (ICB), an immunotherapy increasingly being used in patients with advanced cancers^1–3^. In 2015, the U.S. Food and Drug Administration approved the first oncolytic virus for the treatment of advanced melanoma: Talimogene Laherparepvec (T-VEC) is an engineered herpes simplex virus-1 that allows tumor-selective replication and expresses human granulocyte-macrophage colony-stimulating factor (GM-CSF)^4–6^. Clinical trials on the combination of intratumoral (IT) injection of T-VEC with systemic delivery of ICB agents-- including ipilimumab and pembrolizumab, which blocks the cytotoxic T cell-associated antigen 4 (CTLA-4) and programmed death protein 1 (PD-1), respectively -- have shown enhanced therapeutic efficacy and increased cytotoxic T cell infiltration into tumors^7–9^.

Poxviruses are large cytoplasmic DNA viruses^10^. Preclinical studies and clinical trials have demonstrated the efficacy of using oncolytic vaccinia viruses for the treatment of advanced cancers^11–15^. For example, JX594, also known as Pexastimogene Devacirepvec (Pexa Vec), an oncolytic vaccinia virus featuring the deletion of the thymidine kinase (TK) gene to enhance tumor selectivity and the expression of human GM-CSF, has demonstrated therapeutic efficacy in a Phase II clinical trial for patients with advanced hepatocellular carcinoma (Heo et al., 2012). Unfortunately, phase III clinical trial (NCT02562755) comparing Pexa Vec followed by sorafenib vs. sorafenib alone discontinued enrollment after an interim futility analysis showing lack of efficacy.

While the tumor killing effects of oncolytic viruses have traditionally been attributed to their selective replication within tumor cells, the ability of oncolytic virotherapy to elicit host antitumor immunity plays a crucial role as well^16–20^. However, the mechanisms by which oncolytic virotherapy induces antitumor immunity remain largely unknown. Modified vaccinia virus Ankara (MVA), a highly attenuated vaccinia strain, is an important vaccine vector against various infectious agents^21–26^. We previously reported that MVA infection of conventional dendritic cells (cDCs) triggers type I IFN via the cGAS/STING-mediated cytosolic DNA-sensing pathway^27^. The cGAS/STING and type I IFN pathways are innate sensing mechanisms critical for antiviral and antitumor immunity^28–38^. Infection of cDCs with heat-inactivated MVA (heat-iMVA) generated by heating the virus at 55°C for 1 h leads to higher levels of type I IFN than with MVA. Intratumoral (IT) delivery of heat-iMVA leads to tumor eradication as well as the development of systemic antitumor immunity, which requires CD8^+^ T cells, Batf3-dependent CD103^+^ DCs, as well as the STING-mediated cytosolic DNA-sensing pathway^39^. Our results demonstrate that IT heat-iMVA alters the immunosuppressive tumor microenvironment (TME) by inducing the production of IFN, proinflammatory cytokines, and chemokines both in tumor cells and immune cells via STING, and by activating tumor-infiltrating CD103^+^ DCs that contribute to the priming, expansion and recruitment of activated antitumor CD4^+^ and CD8^+^ T cells and tumor eradication^39^.

In this study, we engineered a replication-competent, oncolytic vaccinia virus (OV-GM) by inserting murine GM-CSF gene into the thymidine kinase (TK) locus of a mutant vaccinia E3LΔ83N (Western Reserve strain), which lacks the Z-DNA-binding domain of vaccinia virulence factor E3. E3LΔ83N replicates efficiently in many cell lines, but is highly attenuated, with reduced virulence of about 1,000-fold compared with wild type vaccinia in *in vivo* infection models^40^. We compared the antitumor effects of IT delivery of live OV vs. live OV-GM vs. heat-inactivated OV-GM (heat-iOV-GM) in bilateral and unilateral murine tumor models in immune-competent syngeneic mice. Expression of murine GM-CSF by live OV-GM slightly improved the generation of effector CD8^+^ and CD4 T^+^ cells in both the injected and non-injected tumors. Although heat-iOV-GM does not express murine GM-CSF, we compared the antitumor effects of heat-iOV-GM with live OV-GM because many oncolytic viral platforms express GM-CSF as a transgene including JX594/Pexa Vec, a clinical oncolytic vaccinia candidate. We found that IT heat-iOV-GM is more effective in eradicating tumors and generating systemic antitumor immunity than live OV-GM in both unilateral and bilateral tumor implantation models. The antitumor effects of both live OV-GM and heat-iOV-GM require Batf3-dependent CD103^+^/CD8α^+^ DCs, which are efficient in cross-presenting tumor antigens. Our results provide strong evidence that viral replication and viral-mediated oncolysis are not required for the generation of antitumor effects by vaccinia virus. Rather, the activation of the host’s immune system -- including the Batf3-dependent CD103^+^ DCs, likely via the cGAS/STING-mediated cytosolic DNA-sensing pathway -- is crucial for the therapeutic efficacy of vaccinia-based immunotherapy. Our findings have important clinical implications for the future design of optimal vaccinia-based cancer immunotherapeutics.

## Methods

### Study design

In this study, we used unilateral and bilateral tumor implantation models to compare the anti-tumor activities of live or heat-inactivated recombinant vaccinia virus expressing murine GM-CSF. We also determined the relative contributions of the cytosolic DNA-sensing pathway and CD103^+^ DCs in the induced antitumor effects using STING^Gt/Gt^ and Batf3^−/−^ mice. In all experiments, animals were assigned to various experimental groups at random. For survival studies, sample sizes of 8-10 mice were used and the experiments were performed two or three times. For experiments designed to evaluate the tumor immune cell infiltrates, 3-5 mice were used for each experiment and the experiments were performed two or three times. For the experiments designed to assess the induction of type I IFN and proinflammatory cytokines and chemokines in tumors, 3-5 mice were used for collecting the tumors and triplicate quantitative real-time PCR analyses were performed for each tumor sample, and the experiments were performed two or three times.

### Viruses and Cell lines

E3LΔ83N virus was kindly provided by Bertram Jacobs (Arizona State University). E3LΔ83N (OV-TK^+^), OV, or OV-GM viruses were propagated in BSC-40 (Africa green monkey kidney cells, ATCC-CRL2761) cells. Viruses were purified through a 36% sucrose cushion. Heat-iOV-GM virus was generated by incubating purified OV-GM virus at 55°C for 1 hour. BSC-40 cells were maintained in DMEM medium containing 5% FBS, penicillin and streptomycin. The murine melanoma B16-F10 cell line was originally obtained from I. Fidler (MD Anderson Cancer Center) and was maintained in RPMI 1640 medium supplemented with 10% FBS, penicillin and streptomycin.

### Mice

Female C57BL/6J mice were purchased from the Jackson Laboratory (Stock # 000664). Batf3^−/−^ mice were from Dr. Kenneth Murphy (Washington University). STING^Gt/Gt^ mice were a kind gift from Dr. Russell Vance (University of California, Berkeley). These mice were maintained in the animal facility at the Sloan Kettering Institute. All procedures were performed in strict accordance with the recommendations in the Guide for the Care and Use of Laboratory Animals of the National Institute of Health. The animal protocol was approved by the Institutional Animal Care and Use Committee at Sloan Kettering Institute.

### Generating recombinant vaccinia virus expressing mGM-CSF

Murine GM-CSF (mGM-CSF) coding sequence was inserted into the pCB vector between Xba I and EcoR I sites. Vaccinia synthetic early and late promoter (PsE/L) was used to express mGM-CSF, and the vaccinia P7.5 promoter was used to express the drug selection gene, the E. coli xanthine-guanine phosphoribosyl transferase gene (gpt). These two expression cassettes were flanked by a partial sequence of the TK gene on each side. To generate recombinant viruses OV (E3LΔ83N-TK^−^) or OV-GM (E3LΔ83N-TK^−^-mGM-CSF), BSC40 cells were seeded into a 6-well plate and were then infected with E3LΔ83N at the multiplicity of infection (MOI) of 0.05. Two hours after virus infection, transfection mixtures containing plasmid DNA and Lipofectamine 3000 (Invitrogen) were added to the well, and the cells were incubated at 37°C for 48 hours. The recombinant viruses were enriched in the gpt selection medium which contained mycophenolic acid (MPA), xanthine and hypoxanthine, and were plaque-purified in the gpt selection medium four times until the respective purified recombinant viruses were obtained. PCR reactions were used to verify the purity of these recombinant viruses. The primer sequences used for the PCR reactions are:

TK-F2: 5’-TGTGAAGACGATAAATTAATGATC-3’;
pCB-R3: 5’-ACCTGATGGATAAAAAGGCG-3’;
TK-F4: 5’-TTGTCATCATGAACGGCGGA-3’;
TK-R4: 5’-TCCTTCGTTTGCCATACGCT-3’;
GM-F: 5’-GGCATTGTGGTCTACAGCCT-3’;
GM-R: 5’-GTGTTTCACAGTCCGTTTCCG-3’;
TK-F5: 5’-GAACGGGACTATGGACGCAT-3’;
TK-R5: 5’-TCGGTTTCCTCACCCAATCG-3’.

### Cytokine assays

Cells were infected with various viruses at a MOI of 10 for 1 h or mock infected. The inoculum was removed and the cells were washed with PBS twice and incubated with fresh medium. Supernatants were collected at various times post infection. Cytokine levels were measured by using enzyme-linked immunosorbent essay (ELISA) kits for murine IFN-α/β (PBL Biomedical Laboratories), IL-6, CCL5, CXCL10, and GM-CSF (R & D systems).

### Western Blot Analysis

B16-F10 cells were infected with OV-GM at a MOI of 10, and cell lysates was collected at different time points after virus infection. Polypeptides were separated by 15% SDS-PAGE, and western blot analysis was performed to determine the expression of mGM-CSF using anti-mGM-CSF antibody (Thermo Fisher). GADPH was used as a loading control.

### mGM-CSF bioactivity assay

B16-F10 cells were infected with OV-GM at a MOI of 10 for 1 hour in a 6-well plate, and the inoculum was removed and cells were washed with PBS. Fresh medium was added to the well, and the culture supernatants were collected at 24 hours after virus infection. The supernatant was UV irradiated and filtered through a 0.2 μm syringe filter (Nalgene). Different dilutions of the supernatants were added to bone marrow cells in RMPI medium. Generation of bone marrow derived dendritic cells (BMDCs) was described previously. After 7 days, cultured DCs were fixed with Fix Buffer I (BD Biosciences) for 15 min at 37°C. Cells were washed, permeabilized with PermBuffer (BD Biosciences) for 30 min on ice, and stained with antibodies against CD11c and CD11b for 30 min. Cells were analyzed using the LSRII Flow cytometer (BD Biosciences) for CD11c^+^ DCs. Data were analyzed with FlowJo software (Treestar).

### Flow cytometry analysis of DC maturation

For DC maturation analysis, BMDCs were generated from C57BL/6J mice and infected with either live OV or live OV-GM at a MOI of 10 or with equivalent amounts of heat-iOV-GM. Cells were collected at 14 h post infection and were then fixed with Fix Buffer I (BD Biosciences) for 15 min at 37°C. Cells were washed, permeabilized with PermBuffer (BD Biosciences) for 30 min on ice, and stained with antibodies against MHC Class I, CD40, CD86, and CD80 for 30 min. Cells were analyzed using the LSRII Flow cytometer (BD Biosciences). Data were analyzed with FlowJo software (Treestar).

### Tumor re-challenge to assess the development of systemic antitumor immunity

The surviving mice (8 weeks after tumor eradication) were re-challenged with intravenous delivery of a lethal dose of B16-F10 (1 × 10^5^ cells in 50 μl PBS) and then euthanized at 3 weeks post re-challenge to evaluate the presence of tumor foci on the surface of lungs

### ELISPOT assay

Spleens were harvested from mice treated with different viruses, and were mashed through a 70 μm strainer (Thermo Fisher Scientific). Red blood cells were lysed using ACK Lysis Buffer (Life Technology) and the cells were re-suspended in RPMI medium. CD8^+^ T cells were then purified using CD8a (Ly-2) MicroBeads from Miltenyi Biotechnology. Enzyme-linked ImmunoSpot (ELISPOT) assay was performed to measure IFN-γ^+^ CD8^+^ T cells according to the manufacturer’s protocol (BD Bioscience). CD8^+^ T cells were mixed with irradiated B16 cells at 1:1 ratio (250,000 cells each) in RPMI medium, and the ELISPOT plate was incubated at 37°C for 16 hours before staining.

### Preparation of single cell suspensions from tumor samples

B16-F10 melanoma cells were implanted intradermally to the right and left flanks of C57BL/6J mice (5 × 10^5^ cells to the right flank and 2.5 × 10^5^ cells to the left flank). PBS, OV, live OV-GM, or heat-iOV-GM viruses (2 × 10^7^ pfu) were injected IT into the tumors on the right flanks 7 days after tumor implantation. The injections were repeated once 3 days later. Tumors were harvested three days after the second injection with forceps and surgical scissors and were weighed. They were then minced prior to incubation with Liberase (1.67 Wünsch U/ml) and DNase (0.2 mg/ml) in serum free RPMI for 30 minutes at 37°C. Cell suspensions were generated by mashing through a 70μm nylon filter, and then washed with complete RPMI.

### Flow cytometry analysis of tumor infiltrating immune cells

Cells were processed for surface labeling with anti-CD3, CD45, CD4, and CD8 antibodies. Live cells are distinguished from dead cells by using fixable dye eFluor506 (eBioscience). They were further permeabilized using permeabilization kit (eBioscience) and stained for Granzyme B. Data were acquired using the LSRII Flow cytometer (BD Biosciences). Data were analyzed with FlowJo software (Treestar).

### RNA isolation and quantitative real-time PCR

B16-F10 melanoma cells were implanted into the right and left flanks of C57BL/6J mice (5 × 10^5^ cells into the right flank and 2.5 × 10^5^ cells into the left flank). PBS, OV, live OV-GM, or heat-iOV-GM viruses were injected IT into the right-side tumors 7 days after tumor implantation. The injection was repeated once 3 days after the first injection. Three days after the second injection, tumors were harvested from euthanized mice with forceps and surgical scissors and minced. RNA was extracted from the tumor lysates with a RNeasy Mini kit (Qiagen) and was reverse transcribed with a First Strand cDNA synthesis kit (Fermentas). Quantitative real-time PCR was performed in triplicate with SYBR Green PCR Mater Mix (Life Technologies) and Applied Biosystems 7500 Real-time PCR Instrument (Life Technologies) using gene-specific primers. Relative expression was normalized to the levels of glyceraldehyde-3-phosphate dehydrogenase (GAPDH). The primer sequences for quantitative real-time PCR are:

Ifnb-F: 5’-TGGAGATGACGGAGAAGATG-3’;
Ifnb-R: 5’-TTGGATGGCAAAGGCAGT-3’;
Il6-F: 5’-AGGCATAACGCACTAGGTTT-3’;
IL6-R: 5’-AGCTGGAGTCACAGAAGGAG-3’;
Ccl4-F: 5’-GCCCTCTCTCTCCTCTTGCT-3’;
Ccl4-R: 5’-CTGGTCTCATAGTAATCCATC-3’;
Ccl5-F: 5’-GCCCACGTCAAGGAGTATTTCTA-3’;
Ccl5-R: 5’-ACACACTTGGCGGTTCCTTC-3’;
Cxcl9-F: 5’-GGAACCCTAGTGATAAGGAATGCA-3’;
Cxcl9-R: 5’-TGAGGTCTTTGAGGGATTTGTAGTG-3’;
Cxcl10-F: 5’-GTCAGGTTGCCTCTGTCTCA-3’;
Cxcl10-R: 5’-TCAGGGAAGAGTCTGGAAAG-3’;
GAPDH-F: 5’-ATCAAGAAGGTGGTGAAGCA-3’;
GAPDH-R: 5’-AGACAACCTGGTCCTCAGTGT-3’

### Unilateral intradermal tumor implantation and intratumoral injection with viruses

All mouse procedures were performed in strict accordance with the recommendations in the Guide for the Care and Use of Laboratory Animals of the National Institute of Health. The protocol was approved by the Committee on the Ethics of Animal Experiments of Sloan-Kettering Cancer Institute.B16-F10 melanoma cells (1× 10^5^ cells in a volume of 50 μl PBS) were implanted intradermally into the shaved skin on the right flank of WT C57BL/6J or Batf3^−/−^ mice. After 7-8 days post implantation, when the tumors reach 3 mm in diameter, they were injected with PBS, live OV-GM (2 × 10^7^ pfu), or with equivalent amounts of heat-iOV-GM when the mice were under anesthesia. Viruses were injected twice weekly. Mice survival was monitored, and tumor sizes were measured twice a week. Tumor volumes were calculated according the following formula: L (length) × W (width) × H (height)/2. Mice were euthanized for signs of distress or when the diameter of the tumor reached 10 mm. Treatments were ended when mice died/euthanized or tumors completely disappeared.

For combination therapy of large tumors, the first injection started when tumor size reaches 5 mm in diameter. Anti-PD-L1 (200 μg per mouse) or isotype control were given intraperitoneally to the mice concurrent with virus treatment throughout the course of study.

### Bilateral tumor implantation model and assessment of therapeutic efficacy of combination therapy with IT injection with viruses plus ICB

B16-F10 melanoma cells were implanted intradermally to the left and right flanks of C57BL/6J mice (5 × 10^5^ to the right flank and 1 × 10^5^ to the left flank). 7-8 days after tumor implantation, when tumor sizes reach 3 mm in diameter at the right flanks, live OV-GM (2 × 10^7^ pfu) or equivalent amounts of heat-iOV-GM were injected IT into the larger tumors on the right flanks. The tumors were injected twice a week concurrently with intraperitoneal delivery of anti-CTLA-4 (100 μg per mouse) or isotype control antibodies. The tumor sizes were measured, and the survival of mice was monitored. Mice were euthanized for signs of distress or when the diameter of the tumor reached 10 mm. Treatments were ended when mice died/euthanized or tumors completely disappeared.

### Statistics

Two-tailed unpaired Student’s *t* test was used for comparisons of two groups in the studies. Survival data were analyzed by log-rank (Mantel-Cox) test. The p values deemed significant are indicated in the figures as follows: *, p < 0.05; **, p < 0.01; ***, p < 0.001; ****, p < 0.0001. The numbers of animals included in the study are discussed in each figure legend.

### Reagents

The commercial sources for reagents were as follows: Antibodies used for flow cytometry were purchased from eBioscience (Live/Dead eFluor 506, CD45.2 Alexa Fluor 700, CD3 PE-Cy7, CD4 Pacific blue-eFluor 450, CD8 PerCP-efluor710, CD11b APC-eFluor 780, MHC Class I APC, CD40 APC, CD80 APC, CD86 APC), Invitrogen (Granzyme B PE-Texas Red), BD Pharmingen (CD11c-PE-Cy7). Murine anti-GM-CSF antibody was purchase from Thermo Fisher. DNAse I and Liberase TL were purchased from Roche. Recombinant murine GM-CSF protein was purchased from GenScript. Therapeutic anti-CTLA4 (clone 9H10 and 9D9) and anti-PD-L1 (clone 10F.9G2) were purchased from BioXcell.

## Results

### Generation of oncolytic vaccinia virus expressing murine granulocyte-macrophage colony-stimulating factor (mGM-CSF)

Oncolytic vaccinia viruses with the deletion of thymidine kinase (TK^−^) are more attenuated and more tumor-selective than TK^+^ viruses^41 42^. Here, we generated a recombinant TK^−^ oncolytic vaccinia virus expressing mGM-CSF under the control of a vaccinia synthetic early/late promoter (PsE/L) (figure. 1A). VACV-E3LΔ83N virus was used as the parental virus (OV-TK^+^). Two recombinant viruses with the loss of part of the TK gene and with and without mGM-CSF (OV and OV-GM) were generated and verified by PCR analyses and sequencing (Supplementary figure. 1A). The replication capacities of OV-TK^+^, OV, and OV-GM in murine B16-F10 cells were determined by infecting them at a MOI of 0.01. OV-TK^+^ replicated efficiently in B16-F10 cells with viral titers increasing by 20,000-fold at 72 h post infection compared with the viral titers at 1 h post infection (figure. 1B). Deletion of the TK gene resulted in a 3-fold decrease in viral replication efficiency in B16-F10 melanoma cells compared with the parental virus. In addition, OV-GM replicated efficiently in murine B16-F10 cells, with a 2800-fold increase of viral titers at 72 h post infection (figure. 1B).

**Figure 1.**
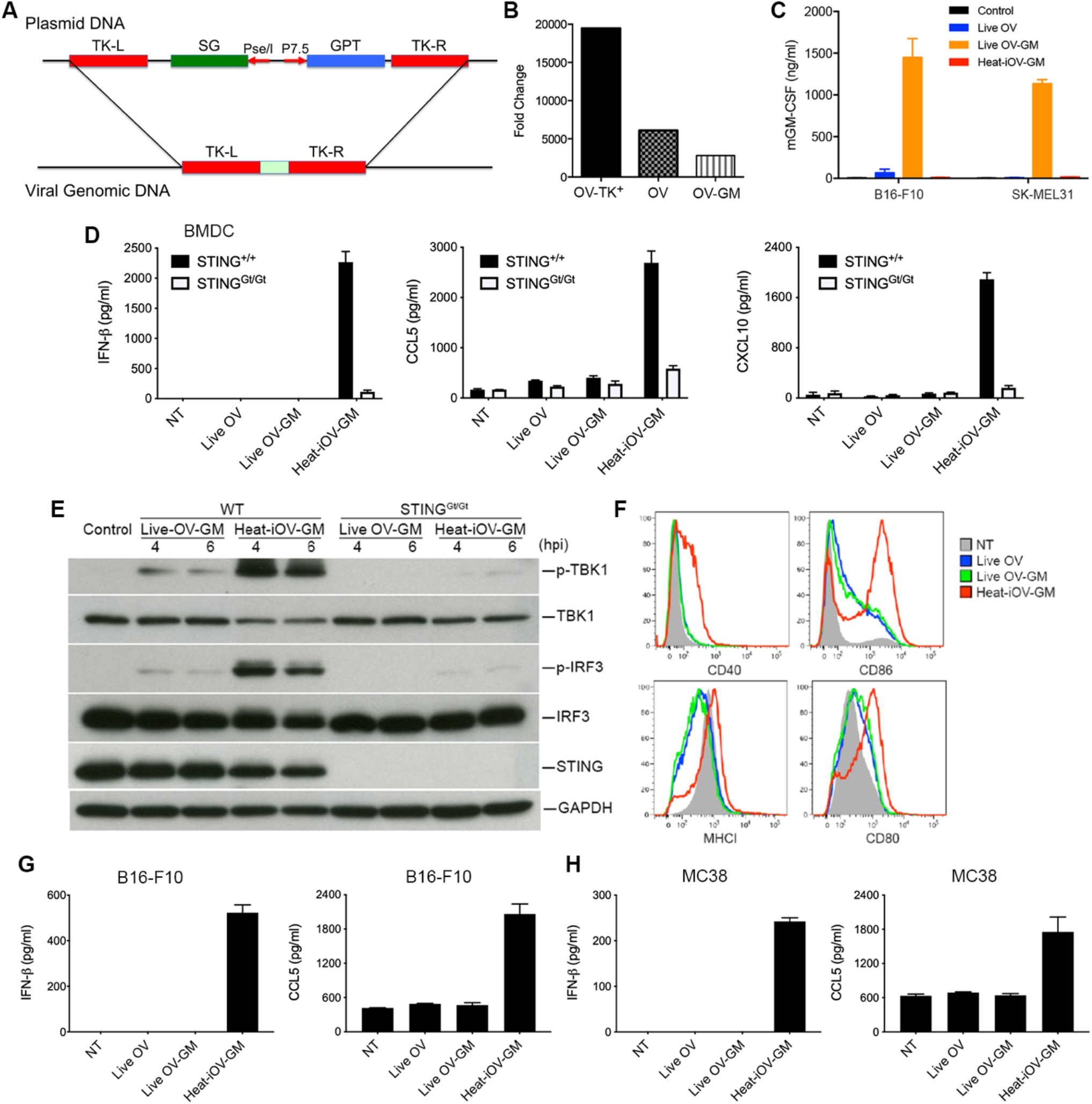
Live oncolytic vaccinia virus fails to induce IFN-β, and CCL5 from infected bone marrow derived dendritic cells (BMDCs), or B16-F10, or MC38 cell. (A) Schematic diagram of homologous recombination between pCB-mGM-CSF plasmid and E3LΔ83N vaccinia viral DNA at the thymidine kinase (TK) locus to generate the recombinant virus E3LΔ83N-TK^−^ (OV) and E3LΔ83N-TK^−^-mGM-CSF (OV-GM). mGM-CSF was expressed under the control of the vaccinia synthetic early and late promoter (PsE/L). (B) Fold changes of viral titers of recombinant viruses in murine B16-F10 melanoma cells at 72 h post infection compared with those at 1 h post infection. B16-F10 melanoma cells were infected with OV-TK+, OV, or OV-GM at a MOI of 0.01. Cells were collected at 1, 24, 48, and 72 h post infection and viral yields (log pfu) were determined by titration on BSC40 cells. (C) mGM-CSF expression of live OV-GM or heat-iOV-GM in B16-F10 or SK-MEL31 cells verified by ELISA. Supernatants were collected at 24 hours post infection. (D) BMDCs were infected with either live OV or live OV-GM at a MOI of 10, or with an equivalent amount of heat-iOV-GM. The supernatants were collected at 22 h post infection. The levels of secreted IFN-β, CCL5 and CXCL10 in the supernatants were determined by ELISA. (E) Western blot analyses of BMDCs from WT or STING^Gt/Gt^ mice infected with either live OV-GM at a MOI of 10 or with an equivalent amount of heat-iOV-GM. The levels of p-TBK1, TBK1, p-IRF3, IRF3, and STING are shown. GAPDH was used as a loading control. hpi: hours post-infection. (F) The expression levels of DC surface markers, MHCI, CD40, CD86, and CD80, on BMDCs infected with either live OV, live OV-GM, or heat-iOV-GM as determined by FACS. NT: no treatment control. (G) The concentrations of secreted IFN-β and CCL5 in the supernatants of murine B16-F10 melanoma cells infected with either live OV or live OV-GM at a MOI of 10, or with an equivalent amount of heat-iOV-GM. (H) The concentrations of secreted IFN-β and CCL5 in the supernatants of murine MC38 colon adenocarcinoma cells infected with either live OV, live OV-GM, or heat-iOV-GM.

To test the expression of mGM-CSF from the OV-GM recombinant viruses, we infected B16-F10 murine melanoma cells with OV-GM at a MOI of 10. Western blot analyses showed the levels of expression of mGM-CSF in both the cell lysates and in the supernatants (Supplementary figure. 1B) at 24 h post infection. The bioactivity of the secreted mGM-CSF was tested by culturing murine bone marrow cells (2.5 × 10^5^) with serial dilution of supernatants obtained from B16-F10 infected with OV-GM (collected at 24 h post infection) or with recombinant mGM-CSF protein (20 ng/ml) for 7 days. The total numbers of CD11c^+^ cells cultured in different conditions are shown (Supplementary figure. 1C). We found that 1:400 dilution of the supernatants collected from OV-GM-infected B16-F10 cells had similar bioactivity to recombinant mGM-CSF (20 ng/ml) (Supplementary figure. 1C). ELISA was used to determine the concentrations of mGM-CSF in the supernatants collected from B16-F10 murine melanoma cells and SK-MEL31 human melanoma cells infected with either live OV or live OV-GM at a MOI of 10, or with equivalent amounts of heat-iOV-GM, at 22 h post infection. The concentrations of mGM-CSF in the supernatants of B16-F10 and SK-MEL31 cells infected with live OV-GM were determined to be 1400 ng/ml and 1200 ng/ml, respectively (figure. 1C). As expected, heat-iOV-GM infection failed to induce mGM-CSF secretion (figure. 1C).

### Heat-inactivated OV-GM (heat-iOV-GM) induces innate immunity in bone marrow-derived dendritic cells (BMDCs) and tumor cells, whereas live OV or live OV-GM does not

We compared the abilities of live OV, live OV-GM, or heat-iOV-GM to induce innate immunity in BMDCs, B16-F10 murine melanoma cells, and MC38 murine colon cancer cells. BMDCs from WT and STING^Gt/Gt^ mice were infected with either live OV or live OV-GM at a MOI of 10, or with equivalent amounts of heat-iOV-GM. Supernatants were collected at 22 h post infection. The concentrations of IFN-β, CCL5, and CXCL10 were determined by ELISA. Whereas live OV or live OV-GM infection failed to induce IFN-β or CXCL10, and only slightly induced CCL5 above background levels, heat-iOV-GM strongly induced IFN-β, CCL5, and CXCL10 in a STING-dependent manner (figure. 1D). Western blot analyses showed that infection of BMDCs with live OV-GM at a MOI of 10 triggered only low levels of phosphorylation of TBK1 and IRF3 at 4 and 6 h post infection, which is dependent on STING. By contrast, infection of BMDCs with heat-iOV-GM strongly induced phosphorylation of TBK1 and IRF3 at 4 and 6 h post infection, which is largely dependent on STING (figure. 1E). FACS analyses of BMDCs infected with either live OV, or live OV-GM, or heat-iOV-GM for 14 h revealed that heat-iOV-GM infection induced the expression level of surface protein levels of MHC class I, CD40, CD86, and CD80 on BMDCs, which are markers of DC maturation. By contrast, live OV or live OV-GM infection resulted in a modest induction of CD86 and CD80 and a reduction of the expression of MHC class I on BMDCs compared with mock treatment control (NT) (figure. 1F). These results indicate that whereas heat-iOV-GM infection of BMDCs induces innate immune responses via the STING-mediated cytosolic DNA-sensing pathway and activates DC maturation, live OV or live OV-GM infection fails to do so.

We also observed that similar findings in murine B16-F10 melanoma and MC38 colon cancer cells infected with either live OV, live OV-GM, or heat-iOV-GM. Heat-iOV-GM infection potently induced IFN-βand CCL5 secretion from B16-F10 (figure. 1G) and MC38 (figure. 1H), but live OV or live OV-GM infection did not.

### The antitumor effects induced by IT live OV-GM are dependent on Batf3-dependent CD103^+^ dendritic cells (DCs)

IT heat-iMVA-induced antitumor effects require Batf3-dependent DCs^39^. Here we used a B16-F10 melanoma unilateral implantation model to test whether live OV-GM also requires Batf3-dependent DCs for antitumor effects. Briefly, B16-F10 melanoma cells (5 × 10^5^ cells) were implanted intradermally into the right flanks of Batf3^−/−^ or wild type (WT) C57BL/6J mice. Seven days after tumor implantation, we injected live OV-GM (2 × 10^7^ pfu) or equivalent amounts of heat-iOV-GM into the tumors on the right flank twice weekly (figure. 2A). We found that IT live OV-GM is effective in delaying tumor growth or eradicating tumors in WT mice, resulting in a 64% survival rate (figure. 2B-C). By contrast, IT live OV-GM is ineffective in Batf3^−/−^ mice, resulting in 0% survival rate. The results are almost indistinguishable from the PBS-treated group with median survival of 17 days in both groups (figure. 2B-C). IT heat-iOV-GM is highly effective in WT mice, resulting in a 92% survival rate, but its efficacy was reduced in Batf3^−/−^ mice, resulting in 0% survival rate. However, there was an extension of median survival from 17 days in the PBS-treated WT mice to 25 days in the heat-iOV-GM-treated Batf3^−/−^ mice (figure. 2B-C). These results are similar to what we reported previously for heat-iMVA^39^. These findings indicate that the antitumor effects of oncolytic DNA virus in a unilateral tumor implantation model require Batf3-dependent CD103^+^ DCs but not viral replication and oncolysis itself. IT heat-iOV-GM is more effective than live OV-GM in eradicating injected tumors, which is likely due to its enhanced ability to induce DC activation and the induction of type I IFN, proinflammatory cytokines, and chemokines in both DCs and tumor cells^39^. Both heat-iOV-GM and heat-iMVA fail to express viral inhibitory proteins that antagonize innate immune sensing mechanisms. We expect that these two inactivated viruses behave similarly: (i) they enter tumor, stromal, and immune cells in the injected tumors, and (ii) viral DNAs gain access to the cytoplasm of infected cells to trigger potent innate immune responses, partly through the activation of the cytosolic DNA-sensing pathway. Therefore, IT heat-iOV-GM leads to the alteration of tumor immunosuppressive microenvironment and enhanced tumor antigen presentation by the CD103^+^ DCs in the tumor-draining lymph nodes (TDLNs).

**Figure 2.**
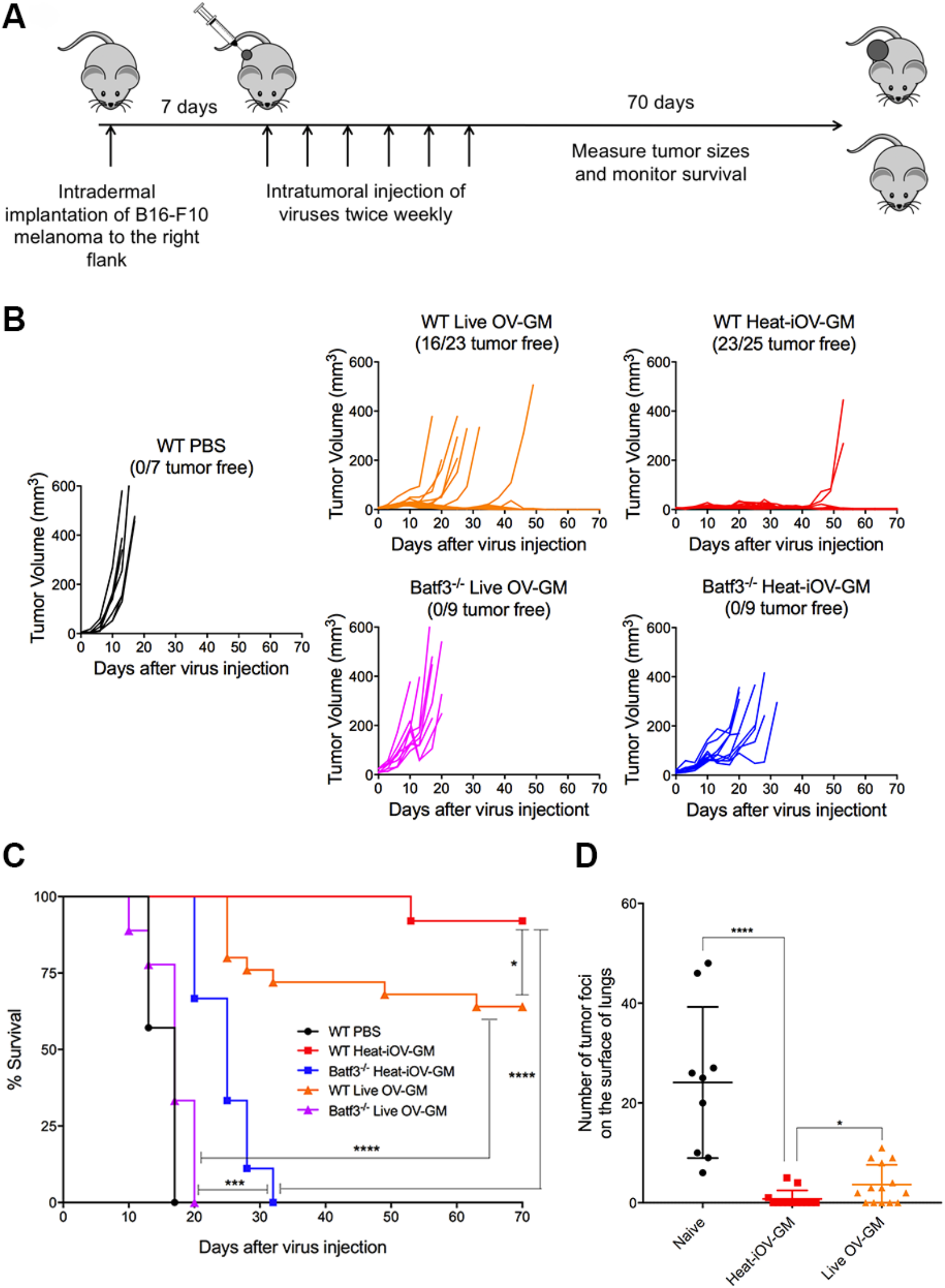
Batf3-dependent CD103+ dendritic cells played an important role in anti-tumor effects of IT live OV-GM and heat-iOV-GM. (A) Tumor implantation and treatment plan in a unilateral B16-F10 intradermal implantation tumor model. (B) Tumor volumes of injected tumors in WT mice treated with either live OV-GM (n=23), heat-iOV-GM (n=25), or PBS control (n=7) or Batf3−/− mice treated with live OV-GM (n=9) or heat-iOV-GM (n=9) over days post treatment. (C) Kaplan-Meier survival curve of WT and Batf3−/− mice treated with PBS, live OV-GM, or heat-iOV-GM. (*P < 0.05; ***P < 0.001; ****P < 0.0001). (D) The number of tumor foci on the surface of lungs collected at 3 weeks from either naïve (n=9), or heat-iOV-GM-treated (n=13), or live OV-GM-treated mice (n=14) after intravenous delivery of 1 × 105 B16-F10 cells (*P < 0.05; ****P < 0.0001, t test).

### IT heat-iOV-GM is more effective in generating long-lasting memory responses against tumor rechallenge in a different organ system compared with IT live OV-GM

IT heat-iMVA-treatment of mice with B16-F10 tumors generates potent systemic antitumor immunity, which results in the rejection of tumor rechallenge through the intravenous (IV) route^39^. Here we compared the efficacy of IT heat-iOV-GM vs. IT live OV-GM in generating systemic antitumor memory responses. IV injection of B16-F10 melanoma cells (1 × 10^5^ cells per mouse) into the surviving mice that were treated previously either with IT heat-iOV-GM or live OV-GM was performed at 8 weeks after the original tumors were eradicated. Mice were euthanized three weeks after rechallenge and the lungs were evaluated under a dissecting microscope for tumor foci on the lung surfaces. Whereas the naïve mice developed an average of 24 tumor foci on the lung surfaces, 5 out of 14 live OV-GM-treated mice failed to develop tumors (with an average of 4 tumor foci on each of the 14 mice), and 10 out of 13 heat-iOV-GM mice rejected tumor challenges (with an average of 0.8 tumor foci on each of the 13 mice) (figure. 2D). These results indicate that IT heat-iOV-GM generated stronger systemic antitumor long-lasting memory immune responses than IT live OV-GM.

### IT heat-iOV-GM induces higher levels of activated CD8+ and CD4+ T cells in the non-injected distant tumors compared with IT live OV-GM

To understand why IT heat-iOV-GM is more effective than live OV-GM in generating antitumor effects, especially in the non-injected distant tumors, we investigated the immune cell infiltrates in both the injected and non-injected tumors in IT heat-iOV-GM or live OV-GM-treated mice. We intradermally implanted 2.5 × 10^5^ B16-F10 melanoma cells to the left flank and 5 × 10^5^ B16-F10 melanoma cells into the right flank of the mice. Seven days post tumor implantation, we injected either 2× 10^7^ pfu of OV, OV-GM, heat-iOV-GM, or PBS into the larger tumors on the right flank. The injection was repeated three days later. Both the injected and non-injected tumors were harvested, and cell suspensions were generated (figure. 3A). We analyzed the live immune cell infiltrates in the tumors by FACS. IT live OV-GM generated higher percentages of Granzyme B^+^ CD8^+^ T cells compared with IT live OV in the distant non-injected tumors (78% in the OV-GM group compared with 56% in the OV group and 54% in the PBS mock-treatment group), although both viruses were highly efficient in the generation of Granzyme B^+^ CD8^+^ T cells in the injected tumors (figure. 3B, 3C). In addition, IT live OV-GM generated higher percentages of Granzyme B^+^ CD4^+^ T cells compared with IT live OV in the distant non-injected tumors (31% in the OV-GM group compared with 16% in the OV group and 13% in the PBS mock-treatment group) (figure. 3D, 3E). In the injected tumors, IT live OV-GM also generated higher percentages of Granzyme B^+^ CD4^+^ T cells compared with IT live OV (96% in the OV-GM group compared with 79% in the OV group and 11% in the PBS mock-treatment group) (figure. 3D, 3E). These results indicate that the expression and secretion of GM-CSF from OV-GM-infected tumor cells have an immune adjuvant effect. However, IT heat-iOV-GM induced higher percentages of Granzyme B^+^ CD8^+^ T cells and Granzyme B^+^ CD4^+^ T cells compared with live OV-GM or live OV in the distant non-injected tumors (94% Granzyme B^+^ CD8^+^ T cells and 62% Granzyme B^+^ CD4^+^ T cells in the heat-iOV-GM group compared with 78% Granzyme B^+^ CD8^+^ T cells and 31% Granzyme B^+^ CD4^+^ T cells in the live OV-GM group) (figure. 3B-E).

**Figure 3.**
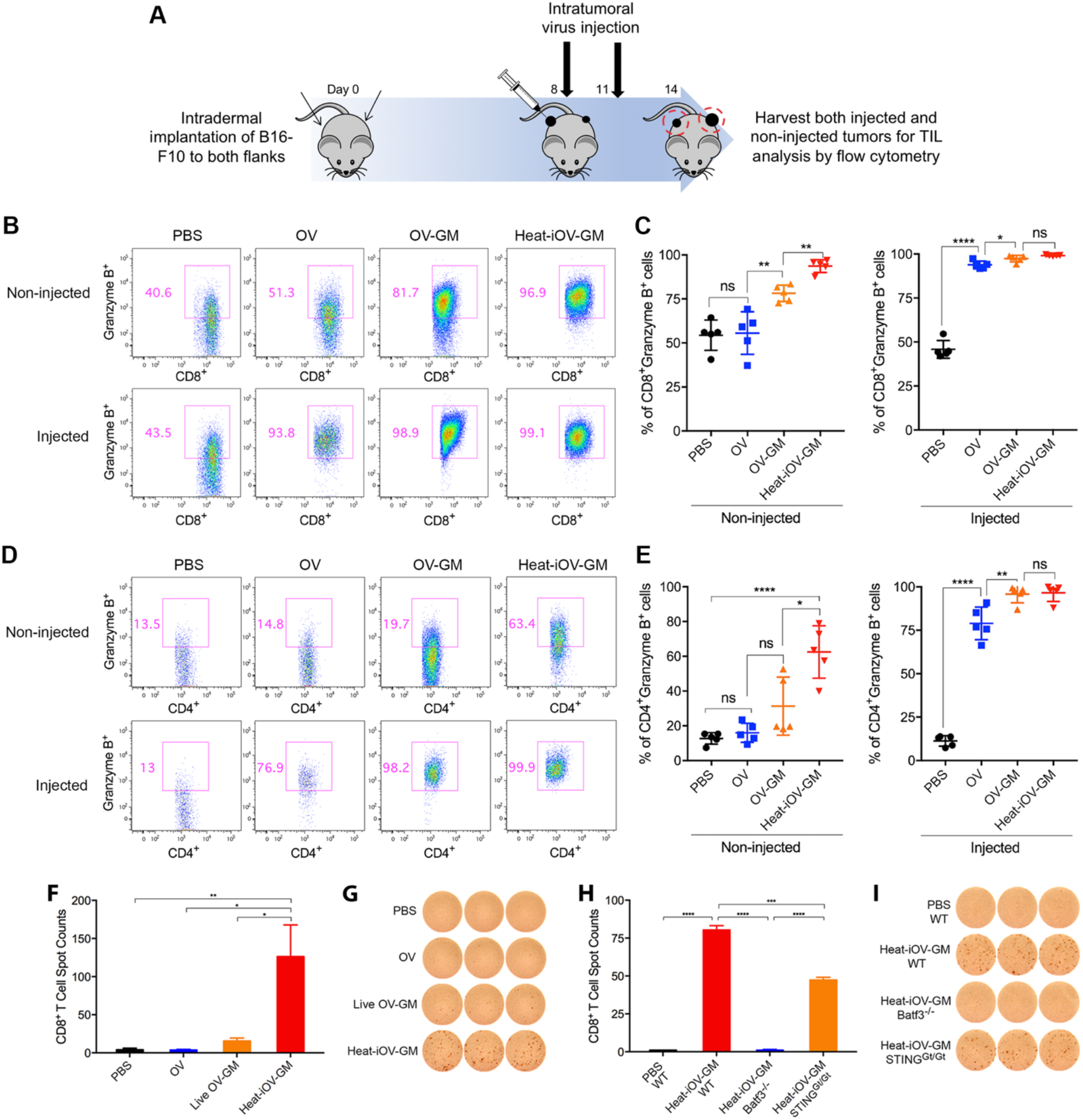
Intratumoral injection of heat-iOV-GM induces higher levels of activated CD8^+^ and CD4^+^ T cells in the non-injected distant tumors. (A) B16-F10 melanoma cells were intradermal implanted to the left and right flanks of mice (2.5 × 10^5^ and 5 × 10^5^ cells, respectively). PBS, OV, live OV-GM, or heat-iOV-GM (2 × 10^7^ pfu) were injected IT into the right-side tumors on day 8 and 11 after tumor implantation. Tumors were harvested 3 days post last virus injection and were used for analyzing the immune cell infiltration by FACS. (B) Representative flow cytometry plot of CD8^+^ T cells expressing Granzyme B in the non-injected or injected tumors from mice treated with PBS, OV, live OV-GM, or heat-iOV-GM. (C) Percentages of CD8^+^ T cells expressing Granzyme B within non-injected and injected tumors. (D) Representative flow cytometry plot of CD4^+^ T cells expressing Granzyme B in the non-injected and injected tumors from mice treated with PBS, OV, live OV-GM, or heat-iOV-GM. (E) Percentages of CD4^+^ T cells expressing Granzyme B within non-injected and injected tumors (n=5, \**P* < 0.05; \*\**P* < 0.01; \*\*\**P* < 0.001; \*\*\*\**P* < 0.0001) (F-I) CD8^+^ T cells from splenocytes of mice treated with different viruses were analyzed for anti-tumor interferon-γ (IFN-γ) activities using ELISPOT assay. (F) IFN-γ^+^ spots per 250,000 purified CD8^+^ T cells from the spleens of the mice treated with IT PBS, OV, live OV-GM, or heat-iOV-GM (n=5, \**P* < 0.05; \*\**P* < 0.01). (G) Representative images from an ELISPOT assay from F. (H) IFN-γ^+^ spots per 250,000 purified CD8^+^ T cells from WT, Batf3^−/−^, or STING^Gt/Gt^ mice treated with IT heat-iOV-GM. (I) ELISPOT images from pooled CD8^+^ T cells of WT, Batf3^−/−^, or STING^Gt/Gt^ mice treated with IT heat-iOV-GM from H. (n=3, \**P* < 0.05; \*\**P* < 0.01; \*\*\**P* < 0.001; \*\*\*\**P* < 0.0001).

### IT heat-iOV-GM induces higher numbers of antitumor CD8+ T cells in the spleens of treated tumor-bearing mice compared with IT live OV-GM

To test whether IT heat-iOV-GM is more effective in generating systemic antitumor immunity compared with IT live OV-GM, we analyzed tumor-specific CD8^+^ T cells in the spleens of tumor-bearing mice treated with either OV, live OV-GM, heat-iOV-GM, or PBS control as described above in a murine B16-F10 bilateral tumor implantation model. Enzyme-linked ImmunoSpot (ELISpot) assay was performed. Briefly, CD8^+^ T cells were isolated from splenocytes and 2.5 × 10^5^ cells were cultured overnight at 37°C in an anti-IFN-γ-coated BD ELISPOT microwells plate. CD8^+^ T Cells were stimulated with B16-F10 cells that were irradiated with an γ-irradiator, and cytokine secretion was detected with an anti-IFN-γ antibody. Whereas CD8^+^ T cells from PBS or OV-treated tumor-bearing mice barely showed any reactivity to B16-F10 cells, CD8^+^ T cells from live OV-GM-treated mice showed some reactivity to B16-F10 cells (figure. 3F-G). By contrast, heat-iOV-GM-treated mice showed much higher numbers of IFN-γ^+^ spots compared with OV, live OV-GM, or PBS-treated mice, with an average of 126 IFN-γ^+^ spots in the heat-iOV-GM group vs. 16 IFN-γ^+^ spots in the live OV-GM group vs. 4 IFN-γ^+^ spots in the OV or PBS group (figure. 3F-G). Similar experiments were performed in MC38 murine colon cancer model and we found that IT heat-iOV-GM generated higher numbers of IFN-γ^+^ spots compared with live OV-GM-treated mice (Supplementary figure. 2A and B). Taken together, these results indicate that IT heat-iOV-GM is more potent compared with live OV-GM in promoting the generation of tumor-specific activated CD8^+^ and CD4^+^ T cells, which are then recruited to inflamed non-injected distant tumors.

### Batf3-dependent CD103^+^ DCs and the STING-mediated cytosolic DNA-sensing pathway are required for the induction of tumor-specific CD8^+^ T cells in the spleens of IT heat-iOV-GM-treated mice

We have previously shown that Batf3-dependent CD103^+^ DCs are critical for the generation of antitumor CD8^+^ T cells in the TDLNs and the recruitment of CD8^+^ and CD4^+^ T cells into injected and non-injected distant tumors in response to IT heat-iMVA^39^. The STING pathway also plays an important role in this process^39^. Here, we tested whether Batf3-dependent CD103^+^ DCs and STING are involved in the generation of tumor-specific CD8+ T cells in the spleens. We found that IT heat-iOV-GM resulted in higher numbers of IFN-γ^+^ spots in WT mice compared with STING^Gt/Gt^ mice, with an average of 80 IFN-γ^+^ spots in the heat-iOV-GM-treated WT mice and 47 IFN-γ^+^ spots in the heat-iOV-GM-treated STING^Gt/Gt^ mice (figure. 3H-I). As expected, IT heat-iOV-GM failed to generate IFN-γ^+^ spots in the Batf3^−/−^ mice (figure. 3H-I). These results further support that IT heat-iOV-GM activates the STING-mediated cytosolic DNA-sensing pathway in Batf3-dependent CD103^+^ DCs to generate systemic antitumor immunity.

### IT heat-iOV-GM induces stronger innate immune responses in the injected tumors compared with live OV-GM

We hypothesized that IT heat-iOV-GM leads to stronger induction of innate immunity in the infected tumor cells and tumor-infiltrating immune cells, compared with IT live OV-GM. To test that, we intradermally implanted B16-F10 melanoma cells into the right flank of C57BL/6J mice; once the tumors were 3-4 mm in diameter, they were injected with either 2 × 10^7^ pfu of live OV-GM or equivalent amounts of heat-iOV-GM. PBS was used as a control. Tumors were harvested one day post infection and mRNAs were extracted. Quantitative real-time PCR analyses of the expression of *Ifnb, Il6, Ccl4, Ccl5, Cxcl9* and *Cxcl10* genes were performed. Whereas IT live OV-GM resulted in modest induction of innate immune responses in the injected tumors compared with IT PBS control, IT heat-iOV-GM resulted in stronger induction of *Ifnb, Il6, Cc4, Ccl5, Cxcl9*, and *Cxcl10* compared with IT live OV-GM (Supplementary figure. 3A-F).

### IT heat-iOV-GM induces higher levels of IFN and proinflammatory cytokines and chemokines in distant non-injected tumors compared with live OV or live OV-GM

Here we compared the innate immunity generated in non-injected distant tumors in mice treated with either IT heat-iOV-GM, live OV-GM, or live OV. Briefly, B16-F10 melanoma cells were implanted intradermally to the left and right flanks of C57BL/6J mice (2.5 × 10^5^ and 5 × 10^5^ cells, respectively). Seven days after tumor implantation, IT injection of 2 × 10^7^ pfu of OV, OV-GM, heat-iOV-GM, or PBS was carried out into the larger tumors on the right flank. The injections were repeated 3 days later. The non-injected tumors on the left flank were harvested 2 days after the last injection, and mRNAs were extracted from the tumor tissue. Quantitative real-time PCR analyses were performed (figure. 4A). IT heat-iOV-GM resulted in the induction of higher levels of *Ifnb, Il6, Ccl4, Ccl5, Cxcl9, and Cxcl10* gene expression in the non-injected distant tumors compared with those mice treated with either live OV-GM, live OV, or PBS control (figure. 4B). These results indicate that IT heat-iOV-GM induces stronger innate immune activation at the non-injected distant tumors compared with IT live OV-GM. Whereas IT live OV is not effective in inducing innate immunity at the non-injected distant tumors compared with PBS mock-treatment control, IT live OV-GM induces slightly higher innate immune responses in the distant non-injected tumors compared with IT live OV (figure. 4B).

**Figure 4.**
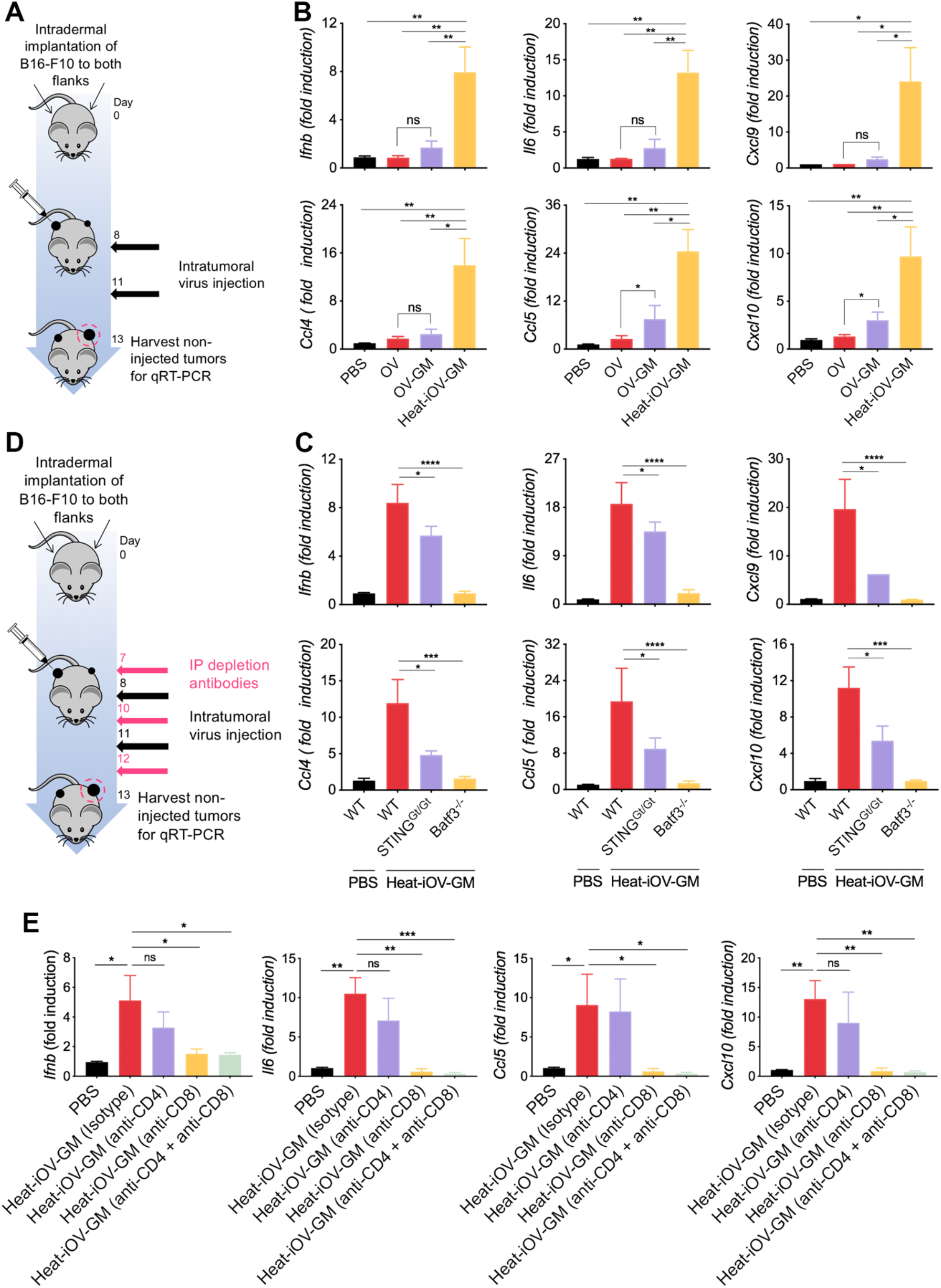
IT heat-iOV-GM induces higher levels of IFN and proinflammatory cytokines and chemokines in distant non-injected tumors than live OV-GM. (A) Tumor implantation and treatment schedule in a bilateral intradermal tumor implantation model. (B) B16-F10 melanoma cells were implanted intradermally on the left and right flanks of C57BL/6J mice. After the tumors are established, the larger tumors on the right flank were injected with either PBS, live OV, live OV-GM, or heat-iOV-GM twice weekly. The non-injected tumors were harvested 2 days after the second injection and RNAs were extracted. Quantitative real-time PCR analyses of *Ifnb, Il6, Ccl4, Ccl5, Cxcl9,* and *Cxcl10* gene expression in non-injected B16-F10 tumors isolated from mice treated with either PBS, OV, live OV-GM, or heat-iOV-GM (n=4-5, \**P* < 0.05, \*\**P* < 0.01, *t* test). (C) The expression of IFN, proinflammatory cytokines and chemokines in non-injected B16-F10 tumors from WT, Batf3^−/−^, or STING^Gt/Gt^ mice treated with heat-iOV-GM were analyzed. Relative expression of *Ifnb, Il6, Ccl4, Ccl5, Cxcl9,* and *Cxcl10* genes was measured by quantitative real-time RT-PCR and was normalized to the expression of GAPDH. Each panel shows the fold changes of the mRNA levels in non-injected tumors from WT, Batf3^−/−^, or STING^Gt/GT^ mice treated with heat-iOV-GM, compared with those from WT mice treated with PBS (n=4, \**P* < 0.05; \*\**P* < 0.01; \*\*\**P* < 0.001; \*\*\*\**P* < 0.0001). (D) Schematic diagram of a bilateral intradermal tumor implantation model with CD4 and/or CD8 depletion. (E) Relative expression level of IFN, proinflammatory cytokines and chemokines in non-injected tumors from each treatment groups were measured by quantitative real-time RT-PCR and was normalized to the expression of GAPDH (n=4, \**P* < 0.05; \*\**P* < 0.01; \*\*\**P* < 0.001; \*\*\*\**P* < 0.0001).

To make sure our observation is not limited to B16-F10 melanoma, we performed similar experiments in a MC38 murine colon cancer model. We confirmed that IT heat-iOV-GM induced higher levels of *Ifnb, Il6, Ccl4, Ccl5, Cxcl9, and Cxcl10* gene expression in both injected tumors (harvested at one day post first injection) and non-injected tumors (harvested two days post second injection) compared with IT live OV-GM (Supplementary figure. 4A-F).

### STING and Batf3-dependent CD103+ DCs contribute to the induction of IFN and proinflammatory cytokines and chemokines by IT heat-iOV-GM in distant non-injected tumors

We hypothesized that the cytosolic DNA-sensing pathway in the Batf3-dependent CD103^+^ DCs might be important for the induction of type I IFN and proinflammatory cytokines and chemokines in response to tumor DNA released from the dying tumor cells. To test that, we intradermally implanted B16-F10 melanoma cells into the left and right flanks of Batf3^−/−^, STING^Gt/Gt^, and WT C57BL/6J mice (2.5 × 10^5^ and 5 × 10^5^ cells, respectively). Seven days after tumor implantation, heat-iOV-GM (an equivalent amount of 2 × 10^7^ pfu of the live virus) or PBS was injected into the larger tumors on the right flank of the mice, with a total of two injections, 3 days apart. The non-injected tumors from the left flank of Batf3^−/−^, STING^Gt/Gt^, and WT mice were harvested at day 3 post the last injection (figure. 4A). Quantitative real-time PCR analyses showed that the induction of *Ifnb*, *Il6*, *Ccl4*, *Ccl5*, *Cxcl9*, and *Cxcl10* gene expression in the non-injected distant tumors of WT mice treated with IT heat-iOV-GM was reduced in STING^Gt/Gt^ mice and abolished in Batf3^−/−^ mice (figure. 4C). These results indicate that STING and Batf3-dependent CD103^+^ DCs play important roles in the induction of IFN and proinflammatory cytokines and chemokines by IT heat-iOV-GM in distant non-injected tumors.

### CD8^+^ T cells are required for the induction of innate immune responses in the distant non-injected tumors

We have previously shown that CD8^+^ T cells are required for heat-iMVA-induced antitumor effects^39^, whereas CD4^+^ T cells are important for the generation of antitumor memory responses. To determine the relative contribution of CD8^+^ and CD4^+^ T cells in mediating innate-immune activation in the non-injected tumors, we either depleted CD8^+^ or CD4^+^ T cells individually or together by administering anti-CD8 and/or CD4 antibodies via intraperitoneal (IP) route one day prior to intratumoral injection of heat-iOV-GM. Two days after the second injection, we isolated the non-injected tumors and performed RT-PCR analyses (figure. 4D). Flow cytometry analyses showed that intratumoral CD4+ and CD8+ T cells were depleted as expected (Supplementary figure. 5). We found that depleting CD8^+^ T cells alone abolished *Ifnb*, *Il6*, *Ccl5*, and *Cxcl10* gene expression in non-injected tumors, whereas depleting CD4^+^ T cells had moderate reduction (figure. 4E). These results indicate that cytotoxic CD8^+^ T cells induced after heat-iOV-GM injection elicit tumor killing in the non-injected tumors and resulting the induction of innate immunity.

### IT heat-iOV-GM generated stronger therapeutic efficacy compared with IT live OV-GM in a B16-F10 bilateral tumor implantation model in the presence or absence of anti-CTLA-4 antibody

Here we investigate the therapeutic efficacy induced by heat-iOV-GM in comparison with live OV-GM in a bilateral tumor implantation model, and whether its combination with systemic delivery of immune checkpoint blockade can further improve the treatment outcome. We implanted B16-F10 cells intradermally into the flanks of C57BL/6J mice, with 5 × 10^5^ to the right flanks and 1 × 10^5^ to the left flanks, and started virus treatment 7-8 days after tumor implantation, when tumor size reaches 3 mm in diameter at the right flanks. Intratumoral injection of either PBS, Live-OV-GM or heat-iOV-GM were given to the tumors on the right flanks twice a week, combined with intraperitoneal (IP) delivery of either anti-CTLA-4 antibody or isotype control. Tumors on the left flanks were not injected with virus. We monitored tumor growth and mice survival (figure 5A). Without anti-CTLA-4 antibody, heat-iOV-GM-treated mice showed improved tumor growth control and survival compared with those treated with live OV-GM, with the extension of median survival from 16.5 days in live OV-GM-treated group to 28 days in heat-iOV-GM treated group (figure 5B and 5C). Combination with immune checkpoint blockade further enhances the abscopal anti-tumor effect induced by heat-iOV-GM. The heat-iOV-GM and anti-CTLA4 combination treatment resulted in a delayed tumor growth and higher rate of tumor regression in the distant tumor compared with the Live-OV-GM and anti-CTLA-4 combination therapy (figure 5C). The cure rate in the heat-iOV-GM plus anti-CTLA4 group was 80%, which is higher than the 40% cure rate in the Live-OV-GM plus anti-CTLA-4 treatment group (figure 5D).

**Figure 5.**
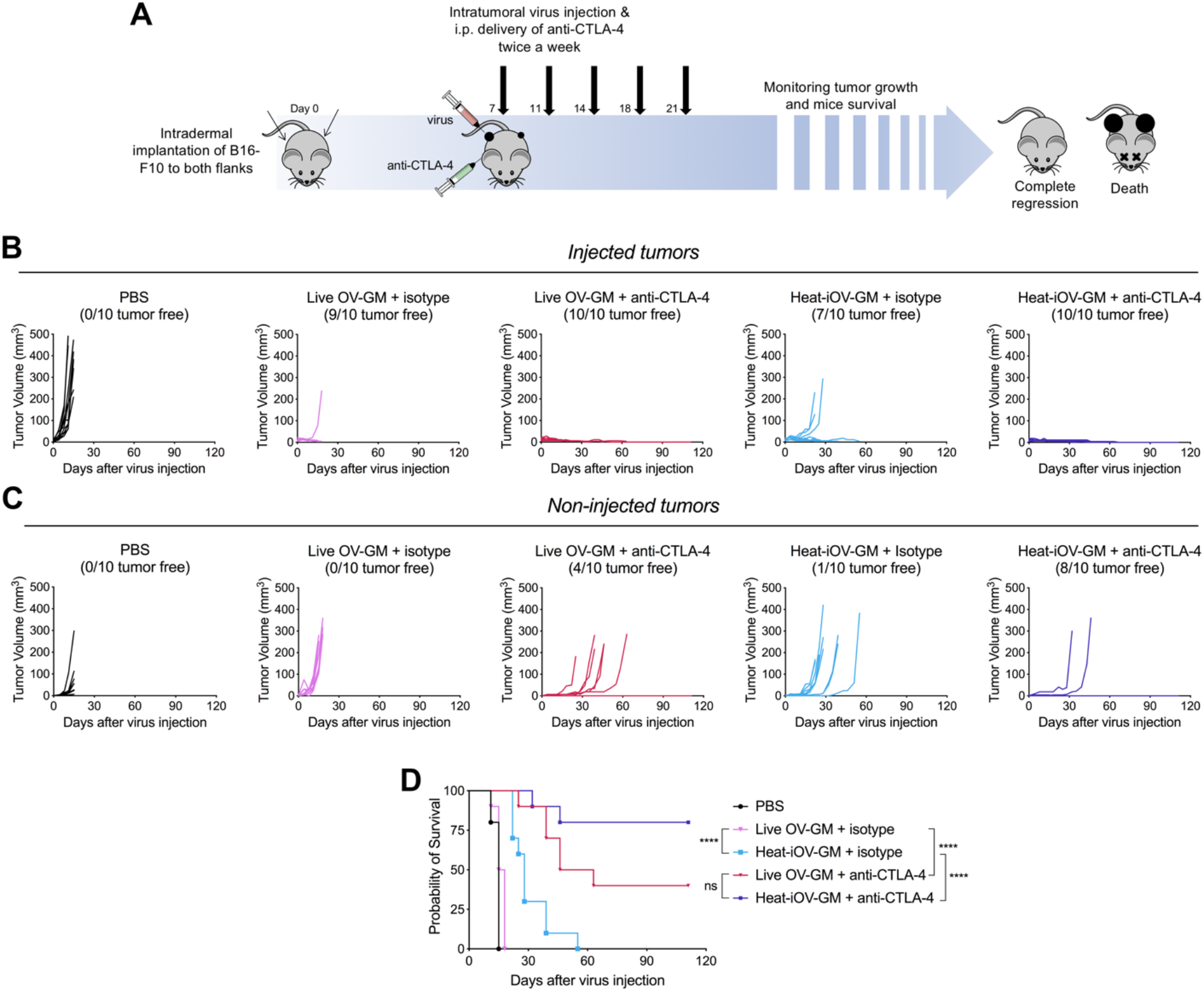
Combination with checkpoint blockade further enhances the abscopal anti-tumor effect induced by heat-iOV-GM. (A) WT C57BL/6J mice were intradermally implanted with B16-F10 tumors on both left and right flanks. Starting from day 7 post implantation, tumor-bearing mice were treated twice weekly with intratumoral injection of live OV-GM or heat-iOV-GM in combination of intraperitoneally injection of isotype control or anti-CTLA-4 (n=10 for all groups). PBS was used as a control. Tumor volume and mice survival were monitored throughout the course of study. (B) Tumor volumes of injected tumors over days of treatment. (C) Tumor volumes of non-injected tumors over days of treatment. (D) Kaplan-Meier survival curve of WT mice treated with PBS, live OV-GM or heat-iOV-GM with or without anti-CTLA-4 (*P < 0.05; ***P < 0.001; ****P < 0.0001).

### Heat-iOV-GM and immune checkpoint blockade combination therapy improves therapeutic efficacy in a large established murine B16-F10 tumor model

We investigated the therapeutic effect of the combination therapy in an aggressive large tumor model. We implanted the B16-F10 cells in the right flank of WT C57BL/6J mice and started virus treatment at a later time point when the tumor size reaches 5 mm in diameter (figure. 6A). While neither the two virus alone nor Live-OV-GM in combination with anti-PD-L1 eradicated the injected tumors, the heat-iOV-GM combined with anti-PD-L1 generated strong antitumor effects leading to tumor regression and elimination in 50% of treated mice (figure. 6B-C). There was an extension of median survival from 14 days in the live-OV-GM plus anti-PD-L1 treated mice to 32 days in the heat-iOV-GM plus anti-PD-L1 treated mice (figure. 6C). These results collectively support that heat-iOV-GM is more immunogenic and generates stronger antitumor effects when combined with ICB compared with live OV-GM plus ICB in both bilateral tumor implantation and large established aggressive tumor models.

**Figure 6.**
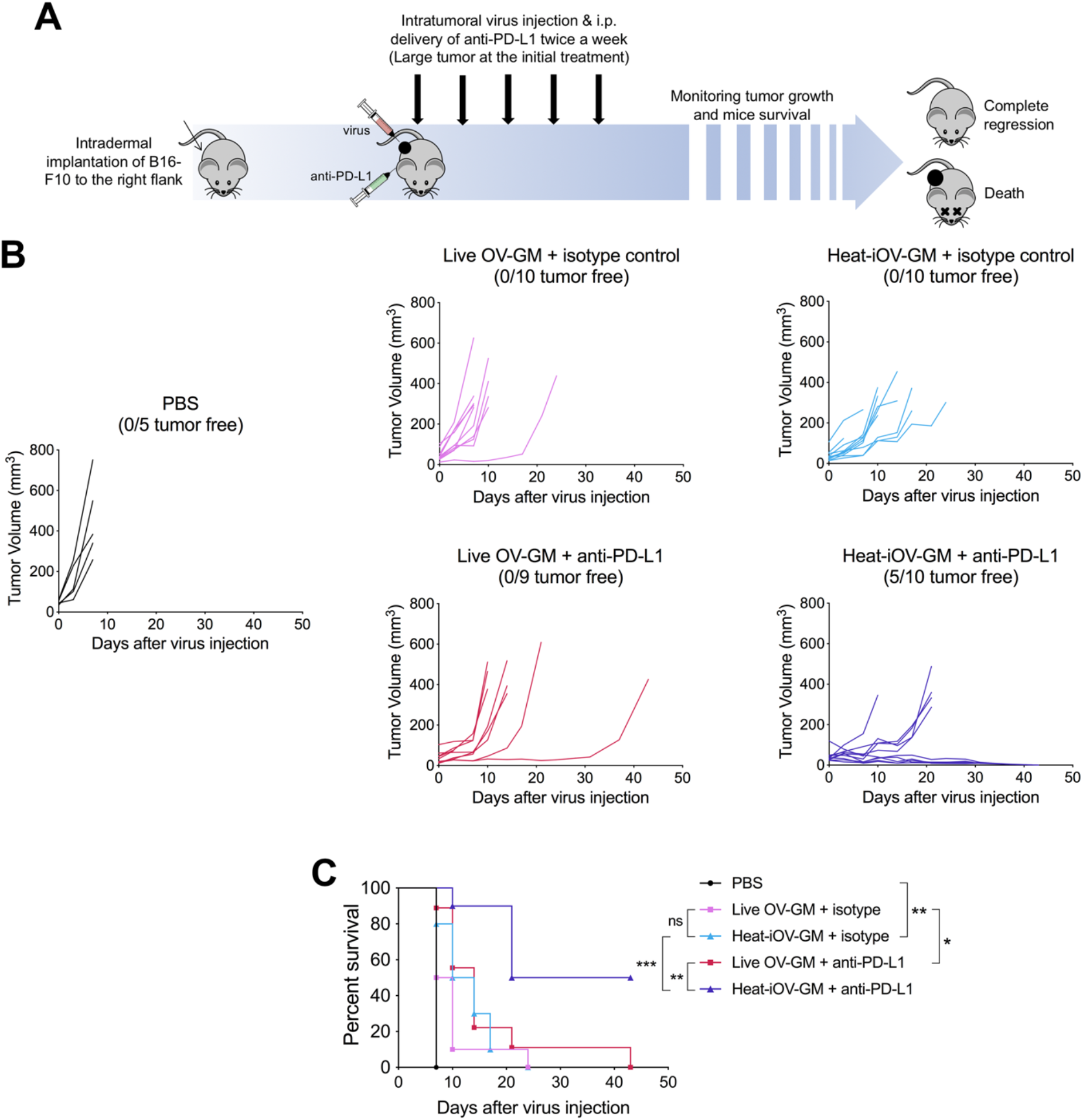
Mice treated with heat-iOV-GM and checkpoint blockade combination therapy show delayed tumor growth and higher survival rate in large tumor model. (A) WT C57BL/6J mice were intradermally implanted with B16-F10 melanoma cells on the right flank. When tumor size reaches 5 mm in diameter, intratumoral injection of live OV-GM or heat-iOV-GM combined with intraperitoneal delivery of anti-PD-L1 or isotype control was initiated and continued twice a week. PBS was used as a control. Tumor growth and mice survival were monitored throughout the course of study. (B) Tumor volumes of injected tumors over days of treatment. (C) Kaplan-Meier survival curve of WT mice treated with PBS, live OV-GM or heat-iOV-GM with or without anti-PD-L1 (n=9 or 10; *P < 0.05; ***P < 0.001; ****P < 0.0001).

## Discussion

Although IT delivery of oncolytic virus Talimogene Laherparepvec (T-VEC) has been approved for the treatment of advanced melanoma as a single agent and IT delivery of T-VEC has been tested in combination with immune checkpoint blockade (ICB) agents in clinical trials for melanoma and other cancers, our understanding of the contribution of viral replication and oncolysis to the generation of antitumor immunity by oncolytic DNA viruses is limited.

In this study, we designed an oncolytic vaccinia virus E3LΔ83N-TK^−^-mGM-CSF (OV-GM) similar to JX594, in which the TK locus was deleted, and the mGM-CSF expression cassette was inserted. JX594 is a leading oncolytic vaccinia virus that has been tested in many clinical trials for various cancers^11–15^. We compared the antitumor immunity of IT live OV-GM vs. IT heat-iOV-GM in both unilateral and bilateral B16-F10 melanoma models, and we found that IT heat-iOV-GM is more effective than IT live OV-GM in eradicating or delaying the growth of both injected and non-injected distant tumors in both models. In the bilateral tumor implantation model, IT heat-iOV-GM induced higher levels of Ifnb, Il6, Ccl4, Ccl5, Cxcl9, and Cxcl10 gene expression in the non-injected distant tumors compared with IT live OV-GM, which correlates with higher numbers of activated CD4^+^ and CD8^+^ T cells in the non-injected tumors in mice treated with IT heat-iOV-GM compared with IT live OV-GM. These results were confirmed in a different murine tumor model MC38 colon cancer, demonstrating that our findings are not limited to one tumor type or microenvironment.

Host type I IFN pathway plays important roles in antitumor immunity^43–45^. Type I IFN signatures correlate with T cell markers in human melanoma metastases^44^. Preclinical studies have shown that IFNAR signaling on dendritic cells, specifically CD103^+^/CD8α^+^ DCs can affect antigen cross-presentation and the generation of antitumor immunity^44 45^. CD103^+^ DCs are tumor-infiltrating DCs, critical for the generation of antitumor immunity, including stimulating naïve and activated CD8^+^ T cells through antigen cross-presentation, and the recruitment of antigen-specific T cells into TME^46 47^. Our *in vitro* and *in vivo* results support our hypothesis that the inferiority of live OV or OV-GM stems from its expression of inhibitory viral genes, which leads to the dampening of type I IFN and pro-inflammatory cytokine and chemokine production in infected bone marrow-derived dendritic cells (BMDCs) and tumor cells. By contrast, heat-inactivated vaccinia failed to express those inhibitory proteins^39 48^. Similar to what we observed with heat-iMVA, infection of heat-inactivated OV-GM in BMDCs and tumor cells leads to the induction of type I IFN, proinflammatory cytokine and chemokine production, whereas live OV or live OV-GM infection of BMDCs, or B16-F10, or MC38 tumor cells fails to induce the above mentioned innate immune mediators. Heat-iOV-GM infection of BMDCs induces DC maturation, whereas live OV or OV-GM infection did not.

Here we propose the following model to explain the induction of innate immunity by heat-iOV-GM in the non-injected distant tumors and the immunological mechanisms underlying the superiority of IT heat-iOV-GM over live OV-GM (Fig. 7). First, compared with Live-OV-GM, heat-iOV-GM infection leads to stronger induction of type I IFN and proinflammatory cytokines and chemokines in the injected tumors via the cGAS/STING-dependent mechanism, which results in stronger activation of CD103^+^ DCs and enhanced tumor-antigen presentation in the TDLNs and spleens. Second, more activated tumor-specific Granzyme B^+^ CD8^+^ and CD4^+^ T cells are then recruited to the distant non-injected tumors to engage in tumor cell killing in heat-iOV-GM-treated mice compared with Live-OV-GM-treated mice. Third, the cGAS/STING-dependent cytosolic-sensing of tumor DNA from dying tumor cells leads to the induction of innate immunity in the non-injected tumors. Finally, heat-iOV-GM treatment generated stronger CD8^+^ T cells-mediated tumor cell killing and higher levels of *Ifnb*, *Il6*, *Ccl4*, *Ccl5*, *Cxcl9* and *Cxcl10* gene expression in the non-injected tumors compared with Live-OV-GM. Based on our findings, we propose that evaluations of both innate and adaptive immunity induced by IT oncolytic viral immunotherapy at non-injected tumors should be included as potential biomarkers for comparing potency and efficacy of various oncolytic constructs in preclinical and clinical studies.

**Figure 7.**
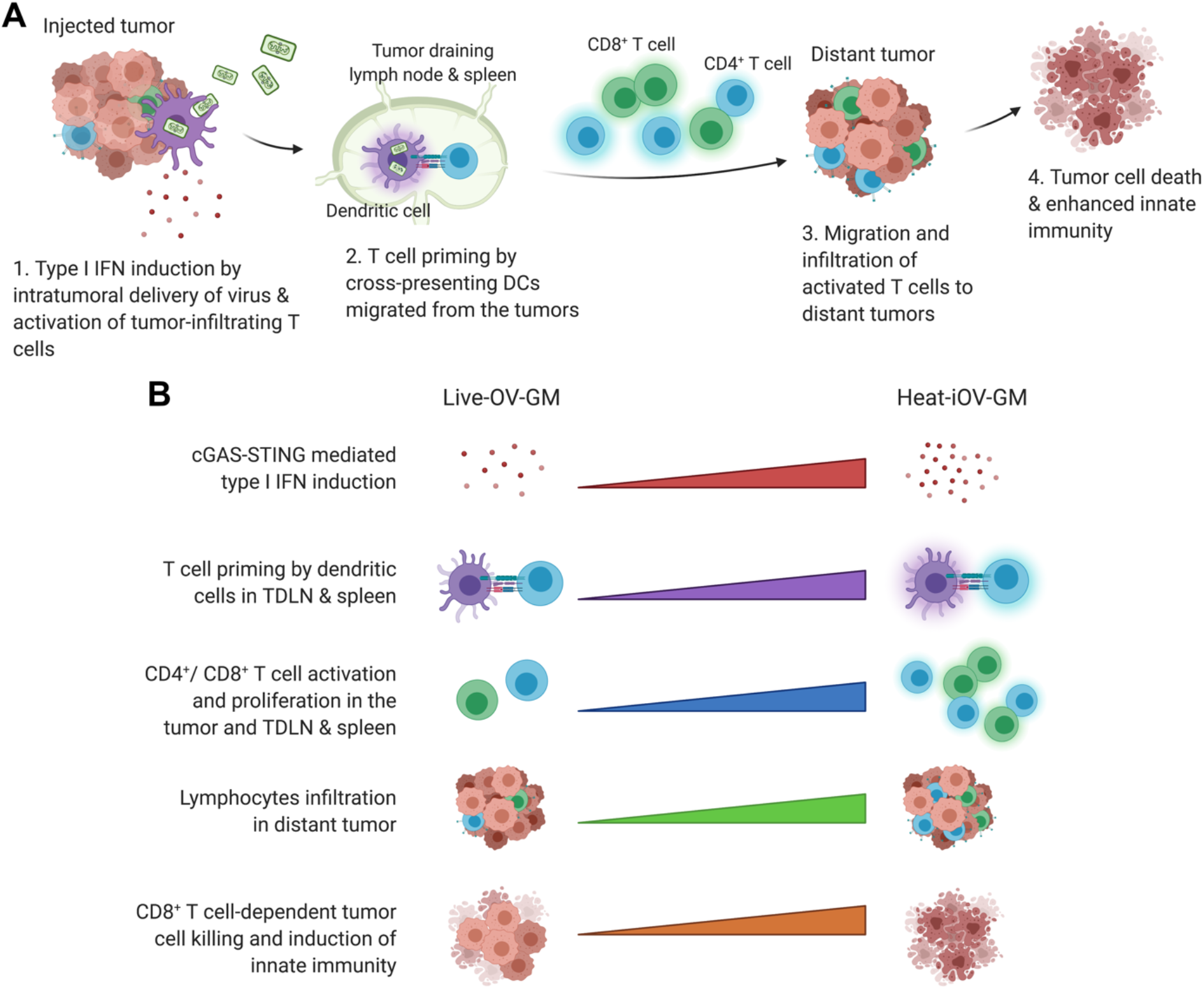
Working model of heat-iOV-GM as a stronger inducer of anti-tumor innate immunity especially in distant tumor compared with live OV-GM. (A) Schematic of induction of innate immunity by intratumoral delivery of heat-iOV-GM vs. live OV-GM in both injected and non-injected distant tumors; created with biorender.com. (B) Comparison of immune activation by heat-iOV-GM vs. live OV-GM. IT delivery of heat-iOV-GM induces (i) higher levels of type I IFN than live OV-GM due to activation of the cGAS/STING-mediated cytosolic DNA-sensing pathway; (ii) stronger T cell priming in TDLN and spleen; and (iii) more activated T cells, which then migrate to the distant tumors resulting in enhanced abscopal tumor cell killing and eventually tumor regression through the induction of a stronger innate immunity.

Our results support that the cGAS/STING-dependent cytosolic-sensing tumor DNA from dying tumor cells in the non-injected tumors leads to the induction of innate immunity. In the absence of STING, IT heat-iOV-GM-induced innate immunity in the non-injected tumors was reduced compared with WT controls, supporting a role of STING in this process. Furthermore, in the absence of Batf3-dependent CD103^+^ DCs, IT heat-iOV-GM-induced innate immunity in the non-injected tumors was abolished. This is consistent with our previous report that in the absence of CD103^+^ DCs, both injected and non-injected tumors failed to recruit anti-tumor CD4^+^ and CD8^+^ T cells in response to IT heat-iMVA treatment^39^. Using ELISpot assay, we showed that Batf3-dependent DCs are crucial for the generation of antitumor CD8^+^ T cells in the spleens of mice after IT heat-iOV-GM treatment. By contrast, IT live OV-GM has limited potency to induce innate immunity at the non-injected distant tumors, which correlates with the lower levels of activated CD8^+^ T cells in the non-injected tumors and spleens compared with those treated with IT heat-iOV-GM. Furthermore, depletion of CD8^+^ T cells from the circulation and tumors abolished IT heat-iOV-GM-induced innate immunity in the non-injected tumors.

Batf3 is a transcription factor that is critical for the development of CD103^+^/CD8α^+^ lineage DCs, which play an important role in cross-presentation of viral and tumor antigens^49 50^. We were surprised by our finding that IT live OV-GM had no antitumor activities in the Batf3^−/−^ mice, whereas IT heat-iOV-GM extended the median survival to 25 days in the Batf3^−/−^ mice compared with 17 days in PBS-treated WT mice. These results suggest that: (i) viral-mediated oncolysis plays little role (if any) in the Batf3^−/−^ mice, which lack CD103^+^/CD8α^+^ DCs; (ii) IT heat-iOV-GM is capable of inducing limited antitumor activity independent of CD103^+^ DCs. This could be related to its ability to induce the production of type I IFN and proinflammatory cytokines and chemokines in other myeloid cells such as CD11b^+^ DCs, plasmacytoid DCs, or tumor-associated macrophages, or inflammatory monocytes, as well as in infected tumors and stromal cells. Further studies to elucidate the contributions of other myeloid cells to heat-iOV-GM-induced antitumor immunity are warranted.

We observed that both IT live OV-GM and IT heat-iOV-GM are capable of generating long-lasting antitumor memory responses through an “in situ vaccination” effect, in which tumor antigens are presented by CD103^+^ DCs to generate tumor-specific CD4^+^ and CD8^+^ T cells in the TDLNs; These cells then return to circulation, are recruited to non-injected distant tumors, or establish residence in secondary lymphoid organs such as the spleen or lymph nodes or in other tissues such as the skin or the lungs. IT heat-iOV-GM is more potent in inducing long-lasting memory responses compared with IT live OV-GM, as 77% of tumor-bearing mice successfully treated with IT heat-iOV-GM rejected tumor rechallenge through IV, whereas only 36% of tumor-bearing mice successfully treated with IT live OV-GM rejected tumor rechallenge. This has important clinical implications because potential viral-based immunotherapy that generates stronger immunological memory will be more effective in preventing cancer recurrence and prolonging patient survival.

In this study, we found heat-iOV-GM performs better than live OV-GM when combined with anti-CTLA-4 antibody in a murine B16-F10 bilateral tumor implantation. The survival advantage of the heat-iOV-GM and anti-CTLA-4 antibody combination is largely due to better control of tumor growth of the non-injected tumors compared with Live-OV-GM plus anti-CTLA-4. This is consistent with the notion that IT heat-iOV-GM generates stronger innate immunity in the non-injected distant tumors compared with Live-OV-GM, which synergizes with systemic delivery of anti-CTLA-4 antibody. In addition, IT heat-iOV-GM plus anti-PD-L1 antibody is more effective in restraining tumor growth compared with IT live OV-GM plus anti-PD-L1 or IT heat-iOV-GM alone in a large established B16-F10 melanoma model. This is likely due to the induction of PD-L1 expression in heat-iOV-GM-infected tumor or immune cells, which can be counteracted by anti-PD-L1 antibody. Together with other published studies, our results support the use of combination therapy of IT immunogenic viruses with systemic delivery of ICB to potentiate antitumor effects in both injected and non-injected tumors^16 39 51^.

## Abbreviations

BMDC: bone marrow-derived dendritic cell
cDCs: conventional dendritic cells
CTLA-4: cytotoxic T cell-associated antigen 4
ELISpot: Enzyme-linked ImmunoSpot
FACS: fluorescence-activated cell sorting
GAPDH: glyceraldehyde-3-phosphate dehydrogenase
gpt: xanthine–guanine phosphoribosyl transferase gene
Heat-iOV-GM: heat-inactivated OV-GM
Heat-iMVA: heat-inactivated MVA
ICB: immune checkpoint blockade
IFN-γ: interferon-γ
IT: Intratumoral
IV: Intravenous
mGM-CSF: murine granulocyte-macrophage colony-stimulating factor
MPA: Mycophenolic acid
MOI: multiplicity of infection
MVA: modified vaccinia virus Ankara
OVs: oncolytic viruses
OV-GM: E3LΔ83N-TK^−^-mGM-CSF
OV: E3LΔ83N-TK^−^-vector
PD-1: programmed cell death protein 1
Pfu: plaque form unit
PsE/L: synthetic early/late promoter
TDLN: tumor draining lymph node
TK: thymidine kinase
TME: tumor microenvironment
T-VEC: Talimogene Laherparepvec
WT: wild type

## Declarations

## Acknowledgements

We thank Dr. Stewart Shuman for critical review of the manuscript. We thank Katharina Shaw for editing. E3LΔ83N virus was kindly provided by Bertram Jacobs (Arizona State University).

## Availability of data and materials

All data published in this report are available on reasonable request.

## Authors’ contributions

W.W. and L.D. designed and performed the experiments, analyzed the data, and prepared the manuscript. P.D. and S.L. performed the experiments, analyzed the data, and assisted in manuscript preparation. N.Y., Y.W., and R.A.G. assisted in some experiments and data interpretation. T.M., J.D.W. assisted in experimental design, data interpretation, and manuscript preparation.

## Patient consent for publication

N/A

## Ethics approval and consent to participate

N/A

## Supplementary Information

This file contains:

- Supplementary Fig. 1
- Supplementary Fig. 2
- Supplementary Fig. 3
- Supplementary Fig. 4
- Supplementary Fig. 5

**Figure S1.**
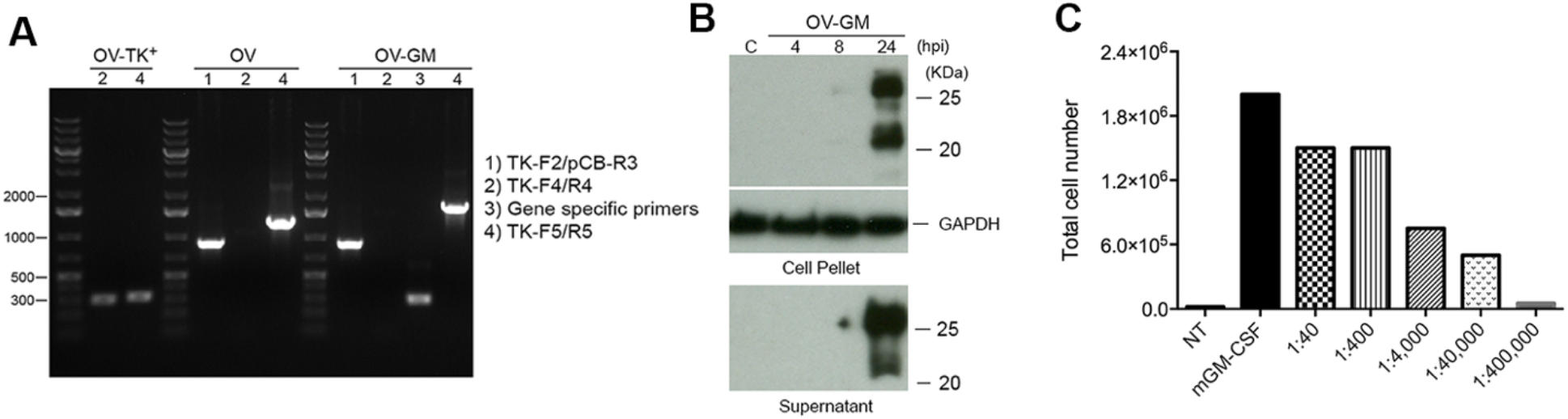
Expression of GM-CSF by B16-F10 cells infected with live OV-GM. (A) PCR verification of recombinant OV and OV-GM. (B) Western blot analysis of mGM-CSF expression in OV-GM-infected murine B16-F10 cells. mGM-CSF protein was detected in both the cell pellet and the culture supernatant. (C) Bioactivity of secreted mGM-CSF protein produced by B16-F10 cells infected with OV-GM. Bone marrow cells (2.5 × 105) from C57BL/6J mice were cultured in the presence of recombinant GM-CSF at 20 ng/ml or with serial dilutions of supernatants from OV-GM-infected B16-F10 melanoma cells for 7 days, and they were subjected to flow cytometry analysis. The total numbers of CD11c+ DCs in various culture conditions are shown.

**Figure S2.**
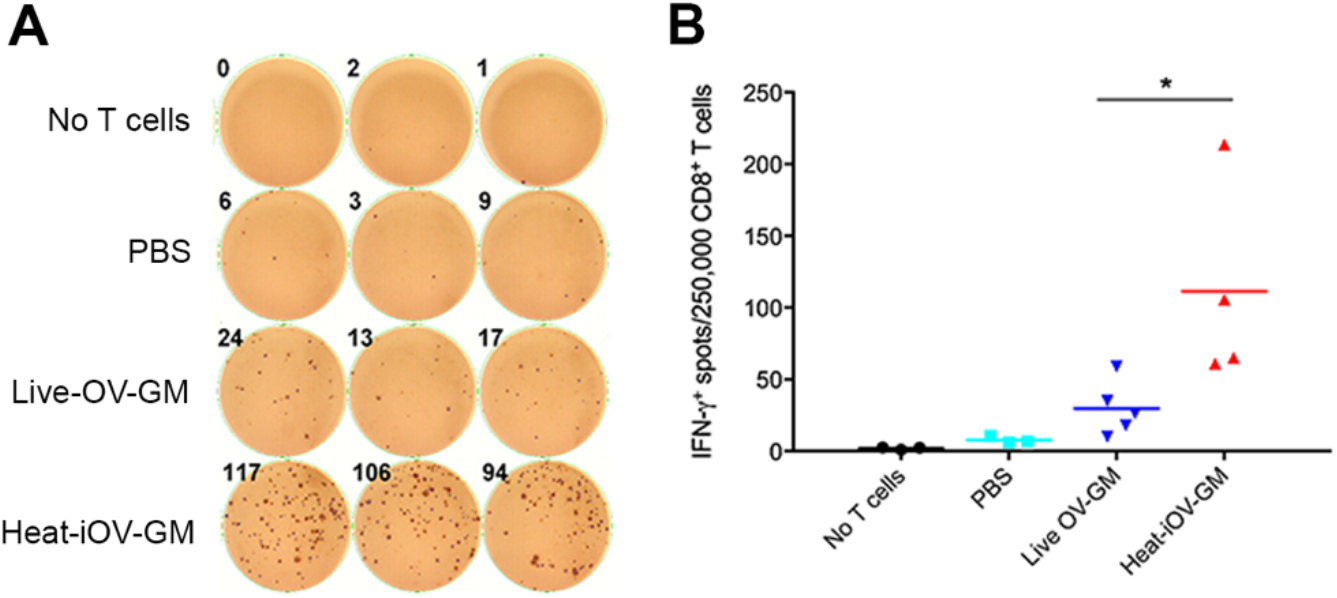
Higher anti-tumor IFN-γ+CD8+ T cells in the spleen from mice treated with heat-iOV-GM compared with live OV-GM in MC38 murine colon cancer model. MC38 tumor cells were intradermally implanted into both flanks of C57BL/6J mice. Established tumors were treated with either Live-OV-GM or heat-iOV-GM. PBS was used as a control. IFN-γ+CD8+ T cells from spleens of MC38 tumor-bearing mice treated with different viruses were analyzed using ELISPOT assay. (A) Representative images from an ELISPOT assay. (B) IFN-γ+ spots per 250,000 purified CD8+ T cells from the spleens of the mice treated with IT PBS, OV, live OV-GM, or heat-iOV-GM (n=5, \**P* < 0.05; \*\**P* < 0.01).

**Figure S3.**
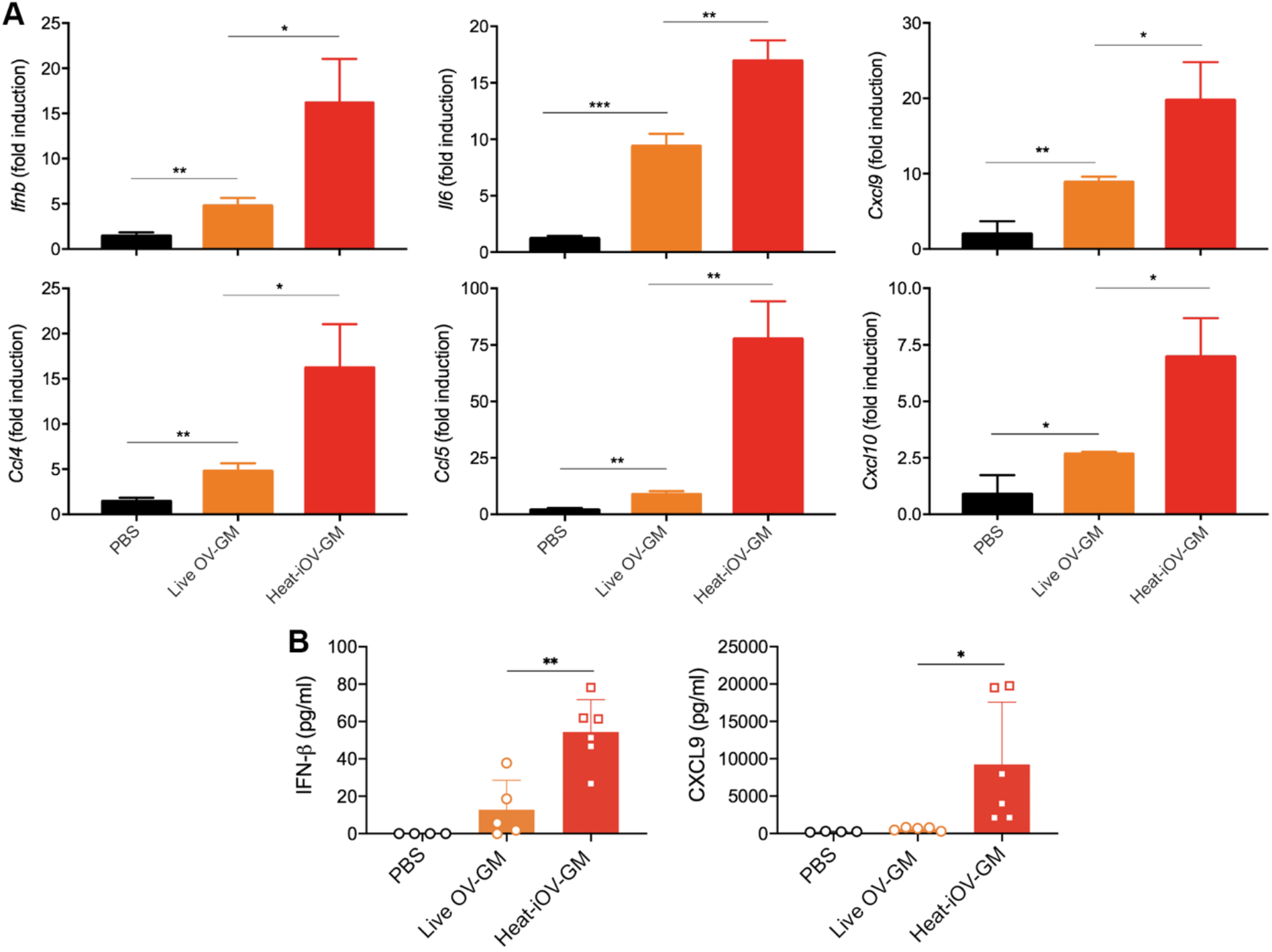
IT heat-iOV-GM induces higher levels of IFN and proinflammatory cytokines and chemokines in the injected tumors compared with IT live OV-GM. B16-F10 melanoma cells were implanted intradermally on the right flank of C57BL/6J mice. Once tumors reach 3-4 mm in diameter, they were injected with either PBS or live OV-GM (2 × 107 pfu), or with equivalent amounts of heat-iOV-GM. The tumors were harvested one day after injection and mRNAs were extracted. (A) Shown here are quantitative real-time PCR analyses of *Ifnb*, *Ccl4*, *Il6*, *Ccl5*, *Cxcl9*, and *Cxcl10* gene expression in the injected B16-F10 tumors from mice treated with either PBS, live OV-GM, or heat-iOV-GM (n=4-5, \**P* < 0.05, \*\**P* < 0.01, \*\*\**P* < 0.001, *t* test). (B) Tumors were harvested one day post injection and homogenized by GentleMACS Dissociator in PBS in the presence of proteinase inhibitor. The levels of IFNβ and CXCL9 were determined by ELISA (n=5-6, \**P* < 0.05, *t* test).

**Figure S4.**
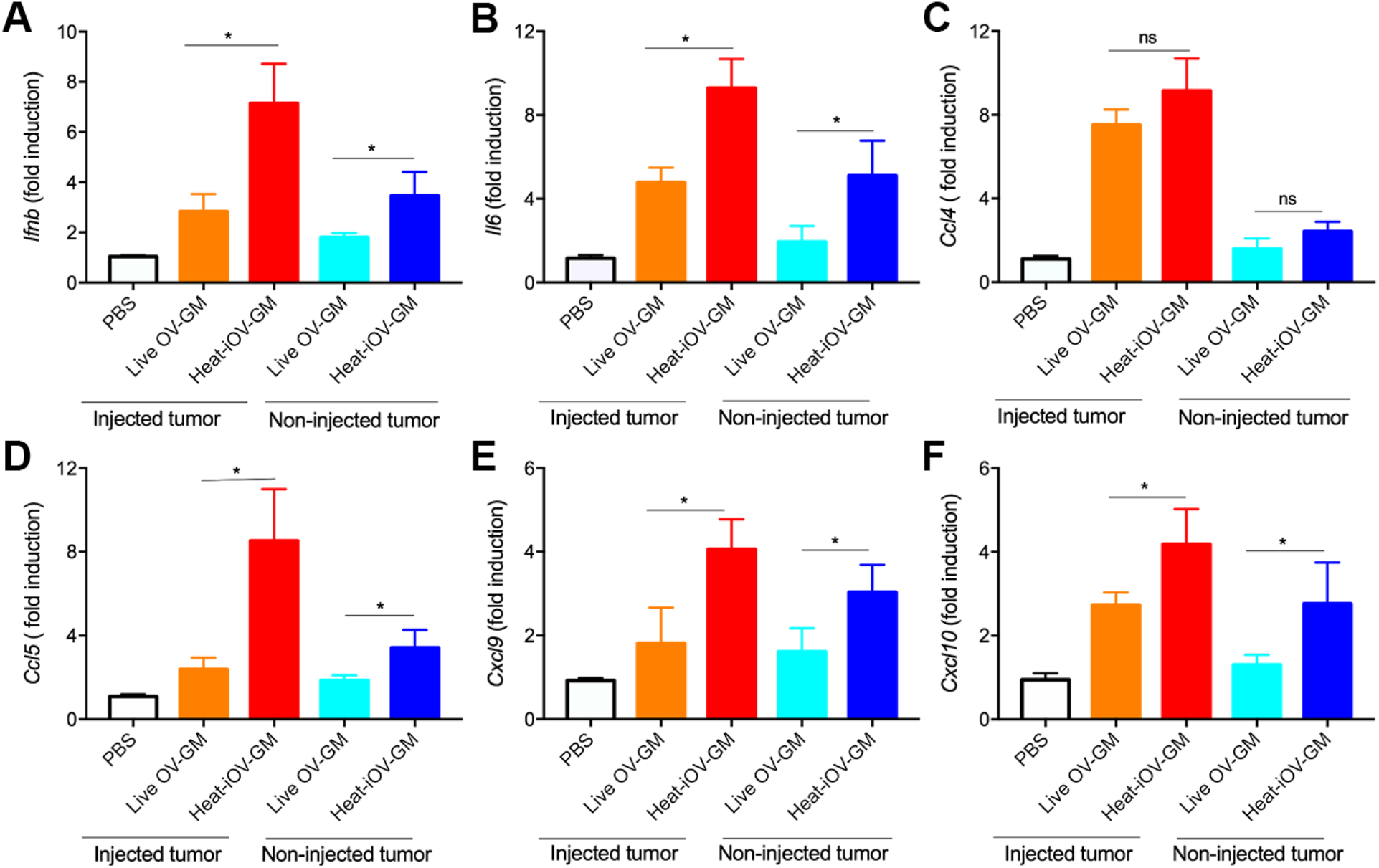
IT heat-iOV-GM induces higher levels of IFN and proinflammatory cytokines and chemokines in both injected and non-injected tumors in MC38 tumor model. MC38 tumor cells were intradermally implanted into both flanks of C57BL/6J mice. Established tumors on the right flanks were treated with either live-OV-GM or heat-iOV-GM. PBS was used as a control. The right flank tumors were harvested one day after first injection. To investigate the innate immunity of the left flank tumors, mice were treated with IT viruses to the right flank tumors twice three days apart. The left flank tumors were harvested one day after the second injection. (A-F) Shown here are quantitative real-time PCR analyses of *Ifnb* (A), *Il6* (B), *Ccl4* (C), *Ccl5* (D), *Cxcl9* (E), and *Cxcl10* (F) gene expression in the injected MC38 tumors from mice treated with either PBS, live OV-GM, or heat-iOV-GM. (n=4-5, \**P* < 0.05, \*\**P* < 0.01, \*\*\**P* < 0.001, *t* test).

**Figure S5.**
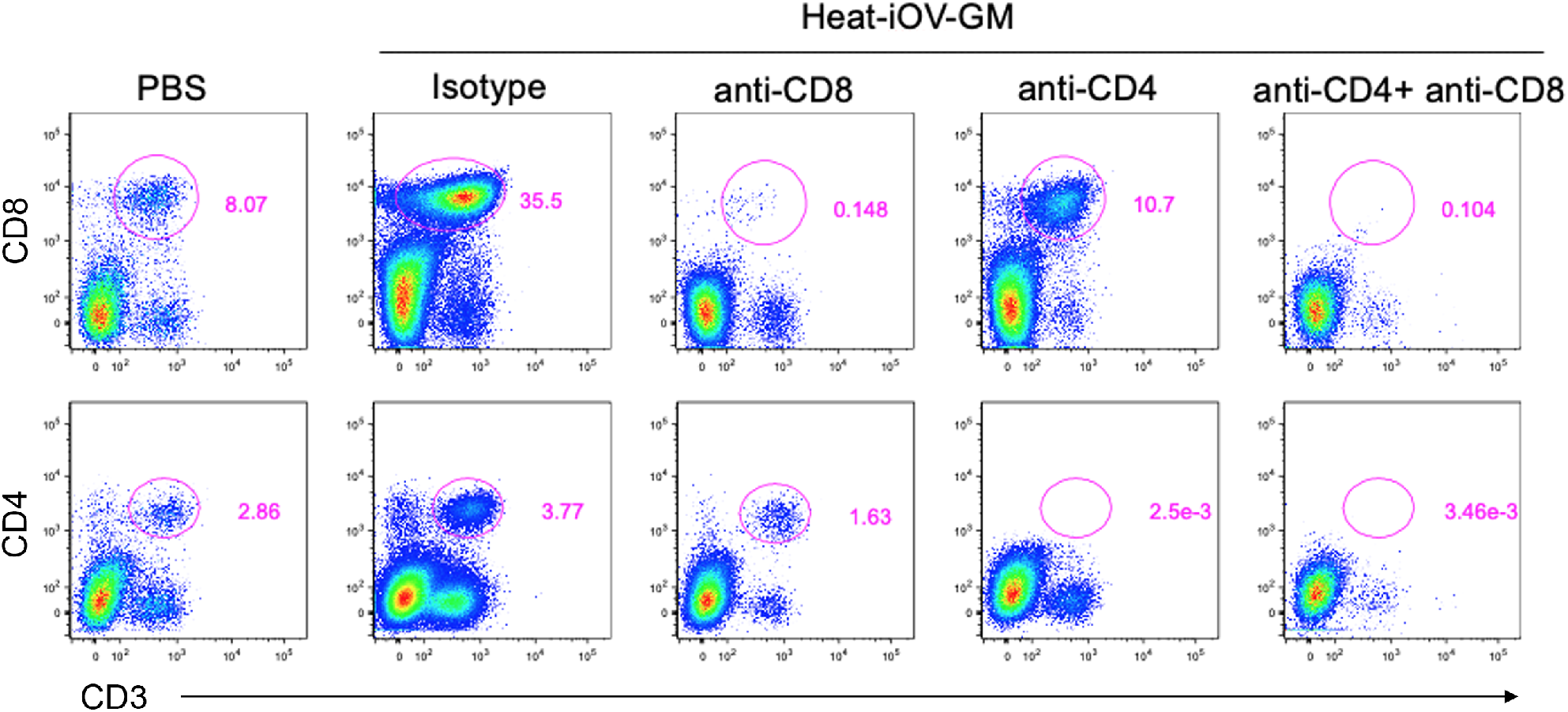
Verification of CD4+ and CD8+ T cell depletion in mice B16-F10 implantation model by flow cytometry. C57BL/6J mice were intradermally implanted with B16-F10 tumors on both the left and right flanks. Established tumors were injected with heat-iOV-GM or PBS control at day 8 and 11 post implantation. Depletion antibody was injected intraperitoneally at day 7, 10, and 12 post implantation. Tumors were harvested at day 13 for TIL analysis. Frequencies of CD4+CD3+ and CD8+CD3+ T cells in tumor samples from each treatment group are shown.

## Reference

1. Russell SJ, Barber GN. Oncolytic Viruses as Antigen-Agnostic Cancer Vaccines. Cancer Cell 2018;33(4):599–605. doi: 10.1016/j.ccell.2018.03.011 [published Online First: 2018/04/11]

2. Bommareddy PK, Shettigar M, Kaufman HL. Integrating oncolytic viruses in combination cancer immunotherapy. Nat Rev Immunol 2018;18(8):498–513. doi: 10.1038/s41577-018-0014-6 [published Online First: 2018/05/11]

3. Zamarin D, Wolchok JD. Potentiation of immunomodulatory antibody therapy with oncolytic viruses for treatment of cancer. Mol Ther Oncolytics 2014;1:14004. doi: 10.1038/mto.2014.4 [published Online First: 2014/01/01]

4. Liu BL, Robinson M, Han ZQ, et al. ICP34.5 deleted herpes simplex virus with enhanced oncolytic, immune stimulating, and anti-tumour properties. Gene therapy 2003;10(4):292–303. doi: 10.1038/sj.gt.3301885

5. Andtbacka RH, Kaufman HL, Collichio F, et al. Talimogene Laherparepvec Improves Durable Response Rate in Patients With Advanced Melanoma. J Clin Oncol 2015;33(25):2780–8. doi: 10.1200/JCO.2014.58.3377

6. Harrington KJ, Andtbacka RH, Collichio F, et al. Efficacy and safety of talimogene laherparepvec versus granulocyte-macrophage colony-stimulating factor in patients with stage IIIB/C and IVM1a melanoma: subanalysis of the Phase III OPTiM trial. Onco Targets Ther 2016;9:7081–93. doi: 10.2147/OTT.S115245

7. Puzanov I, Milhem MM, Minor D, et al. Talimogene Laherparepvec in Combination With Ipilimumab in Previously Untreated, Unresectable Stage IIIB-IV Melanoma. J Clin Oncol 2016;34(22):2619–26. doi: 10.1200/JCO.2016.67.1529

8. Chesney J, Puzanov I, Collichio F, et al. Randomized, Open-Label Phase II Study Evaluating the Efficacy and Safety of Talimogene Laherparepvec in Combination With Ipilimumab Versus Ipilimumab Alone in Patients With Advanced, Unresectable Melanoma. J Clin Oncol 2017:JCO2017737379. doi: 10.1200/JCO.2017.73.7379

9. Ribas A, Dummer R, Puzanov I, et al. Oncolytic Virotherapy Promotes Intratumoral T Cell Infiltration and Improves Anti-PD-1 Immunotherapy. Cell 2017;170(6):1109–19 e10. doi: 10.1016/j.cell.2017.08.027

10. Moss B. Poxviridae: The viruses and their replication. : Lippincott Williams & Wilkins 2007.

11. Park BH, Hwang T, Liu TC, et al. Use of a targeted oncolytic poxvirus, JX-594, in patients with refractory primary or metastatic liver cancer: a phase I trial. Lancet Oncol 2008;9(6):533–42. doi: S1470-2045(08)70107-4 [pii] 10.1016/S1470-2045(08)70107-4 [published Online First: 2008/05/23]

12. Liu TC, Hwang T, Park BH, et al. The targeted oncolytic poxvirus JX-594 demonstrates antitumoral, antivascular, and anti-HBV activities in patients with hepatocellular carcinoma. Mol Ther 2008;16(9):1637–42. doi: mt2008143 [pii] 10.1038/mt.2008.143 [published Online First: 2008/07/17]

13. Breitbach CJ, Burke J, Jonker D, et al. Intravenous delivery of a multi-mechanistic cancer-targeted oncolytic poxvirus in humans. Nature 2011;477(7362):99–102. doi: 10.1038/nature10358 nature10358 [pii] [published Online First: 2011/09/03]

14. Parato KA, Breitbach CJ, Le Boeuf F, et al. The oncolytic poxvirus JX-594 selectively replicates in and destroys cancer cells driven by genetic pathways commonly activated in cancers. Mol Ther 2012;20(4):749–58. doi: 10.1038/mt.2011.276 [published Online First: 2011/12/22]

15. Heo J, Reid T, Ruo L, et al. Randomized dose-finding clinical trial of oncolytic immunotherapeutic vaccinia JX-594 in liver cancer. Nat Med 2013;19(3):329–36. doi: 10.1038/nm.3089 [published Online First: 2013/02/12]

16. Zamarin D, Holmgaard RB, Subudhi SK, et al. Localized oncolytic virotherapy overcomes systemic tumor resistance to immune checkpoint blockade immunotherapy. Sci Transl Med 2014;6(226):226ra32. doi: 10.1126/scitranslmed.3008095 [published Online First: 2014/03/07]

17. Bell J, McFadden G. Viruses for tumor therapy. Cell host & microbe 2014;15(3):260–5. doi: 10.1016/j.chom.2014.01.002

18. Kaufman HL, Kohlhapp FJ, Zloza A. Oncolytic viruses: a new class of immunotherapy drugs. Nat Rev Drug Discov 2015;14(9):642–62. doi: 10.1038/nrd4663

19. Lemay CG, Keller BA, Edge RE, et al. Oncolytic Viruses: The Best is Yet to Come. Curr Cancer Drug Targets 2017 doi: 10.2174/1568009617666170206111609

20. Davola ME, Mossman KL. Oncolytic viruses: how “lytic” must they be for therapeutic efficacy? Oncoimmunology 2019;8(6):e1581528. doi: 10.1080/2162402X.2019.1596006 [published Online First: 2019/05/10]

21. Sutter G, Staib C. Vaccinia vectors as candidate vaccines: the development of modified vaccinia virus Ankara for antigen delivery. Current drug targets Infectious disorders 2003;3(3):263–71. [published Online First: 2003/10/08]

22. McCurdy LH, Larkin BD, Martin JE, et al. Modified vaccinia Ankara: potential as an alternative smallpox vaccine. Clin Infect Dis 2004;38(12):1749–53. doi: 10.1086/421266 [published Online First: 2004/07/01]

23. Vollmar J, Arndtz N, Eckl KM, et al. Safety and immunogenicity of IMVAMUNE, a promising candidate as a third generation smallpox vaccine. Vaccine 2006;24(12):2065–70. doi: S0264-410X(05)01165-5 [pii] 10.1016/j.vaccine.2005.11.022 [published Online First: 2005/12/13]

24. Gomez CE, Najera JL, Krupa M, et al. The poxvirus vectors MVA and NYVAC as gene delivery systems for vaccination against infectious diseases and cancer. Curr Gene Ther 2008;8(2):97–120. [published Online First: 2008/04/09]

25. Goepfert PA, Elizaga ML, Sato A, et al. Phase 1 safety and immunogenicity testing of DNA and recombinant modified vaccinia Ankara vaccines expressing HIV-1 virus-like particles. J Infect Dis 2011;203(5):610–9. doi: jiq105 [pii] 10.1093/infdis/jiq105 [published Online First: 2011/02/02]

26. Gomez CE, Najera JL, Krupa M, et al. MVA and NYVAC as vaccines against emergent infectious diseases and cancer. Curr Gene Ther 2011;11(3):189–217. [published Online First: 2011/04/02]

27. Dai P, Wang W, Cao H, et al. Modified vaccinia virus Ankara triggers type I IFN production in murine conventional dendritic cells via a cGAS/STING-mediated cytosolic DNA-sensing pathway. PLoS Pathog 2014;10(4):e1003989. doi: 10.1371/journal.ppat.1003989 [published Online First: 2014/04/20]

28. Ishikawa H, Ma Z, Barber GN. STING regulates intracellular DNA-mediated, type I interferon-dependent innate immunity. Nature 2009;461(7265):788–92. doi: nature08476 [pii] 10.1038/nature08476 [published Online First: 2009/09/25]

29. Sun L, Wu J, Du F, et al. Cyclic GMP-AMP synthase is a cytosolic DNA sensor that activates the type I interferon pathway. Science 2013;339(6121):786–91. doi: 10.1126/science.1232458 [published Online First: 2012/12/22]

30. Wu J, Sun L, Chen X, et al. Cyclic GMP-AMP is an endogenous second messenger in innate immune signaling by cytosolic DNA. Science 2013;339(6121):826–30. doi: 10.1126/science.1229963 [published Online First: 2012/12/22]

31. Li X, Shu C, Yi G, et al. Cyclic GMP-AMP synthase is activated by double-stranded DNA-induced oligomerization. Immunity 2013;39(6):1019–31. doi: 10.1016/j.immuni.2013.10.019 [published Online First: 2013/12/18]

32. Gao P, Ascano M, Wu Y, et al. Cyclic [G(2′,5′)pA(3′,5′)p] is the metazoan second messenger produced by DNA-activated cyclic GMP-AMP synthase. Cell 2013;153(5):1094–107. doi: 10.1016/j.cell.2013.04.046 [published Online First: 2013/05/08]

33. Gao P, Ascano M, Zillinger T, et al. Structure-function analysis of STING activation by c[G(2′,5′)pA(3′,5′)p] and targeting by antiviral DMXAA. Cell 2013;154(4):748–62. doi: 10.1016/j.cell.2013.07.023 [published Online First: 2013/08/06]

34. Civril F, Deimling T, de Oliveira Mann CC, et al. Structural mechanism of cytosolic DNA sensing by cGAS. Nature 2013;498(7454):332–7. doi: 10.1038/nature12305 [published Online First: 2013/06/01]

35. Ablasser A, Goldeck M, Cavlar T, et al. cGAS produces a 2′-5′-linked cyclic dinucleotide second messenger that activates STING. Nature 2013;498(7454):380–4. doi: 10.1038/nature12306 [published Online First: 2013/06/01]

36. Diner EJ, Burdette DL, Wilson SC, et al. The innate immune DNA sensor cGAS produces a noncanonical cyclic dinucleotide that activates human STING. Cell reports 2013;3(5):1355–61. doi: 10.1016/j.celrep.2013.05.009 [published Online First: 2013/05/28]

37. Woo SR, Fuertes MB, Corrales L, et al. STING-dependent cytosolic DNA sensing mediates innate immune recognition of immunogenic tumors. Immunity 2014;41(5):830–42. doi: 10.1016/j.immuni.2014.10.017

38. Deng L, Liang H, Xu M, et al. STING-Dependent Cytosolic DNA Sensing Promotes Radiation-Induced Type I Interferon-Dependent Antitumor Immunity in Immunogenic Tumors. Immunity 2014;41(5):843–52. doi: 10.1016/j.immuni.2014.10.019

39. Dai P, Wang W, Yang N, et al. Intratumoral delivery of inactivated modified vaccinia virus Ankara (iMVA) induces systemic antitumor immunity via STING and Batf3-dependent dendritic cells. Sci Immunol 2017;2(11) doi: 10.1126/sciimmunol.aal1713 [published Online First: 2017/08/02]

40. Brandt TA, Jacobs BL. Both carboxy- and amino-terminal domains of the vaccinia virus interferon resistance gene, E3L, are required for pathogenesis in a mouse model. J Virol 2001;75(2):850–6. doi: 10.1128/JVI.75.2.850-856.2001 [published Online First: 2001/01/03]

41. Buller RM, Chakrabarti S, Cooper JA, et al. Deletion of the vaccinia virus growth factor gene reduces virus virulence. J Virol 1988;62(3):866–74. [published Online First: 1988/03/01]

42. Puhlmann M, Brown CK, Gnant M, et al. Vaccinia as a vector for tumor-directed gene therapy: biodistribution of a thymidine kinase-deleted mutant. Cancer Gene Ther 2000;7(1):66–73. doi: 10.1038/sj.cgt.7700075

43. Dunn GP, Bruce AT, Sheehan KC, et al. A critical function for type I interferons in cancer immunoediting. Nat Immunol 2005;6(7):722–9. doi: 10.1038/ni1213

44. Fuertes MB, Kacha AK, Kline J, et al. Host type I IFN signals are required for antitumor CD8+ T cell responses through CD8{alpha}+ dendritic cells. J Exp Med 2011;208(10):2005–16. doi: 10.1084/jem.20101159 [published Online First: 2011/09/21]

45. Diamond MS, Kinder M, Matsushita H, et al. Type I interferon is selectively required by dendritic cells for immune rejection of tumors. J Exp Med 2011;208(10):1989–2003. doi: 10.1084/jem.20101158 [published Online First: 2011/09/21]

46. Broz ML, Binnewies M, Boldajipour B, et al. Dissecting the tumor myeloid compartment reveals rare activating antigen-presenting cells critical for T cell immunity. Cancer Cell 2014;26(5):638–52. doi: 10.1016/j.ccell.2014.09.007

47. Spranger S, Dai D, Horton B, et al. Tumor-Residing Batf3 Dendritic Cells Are Required for Effector T Cell Trafficking and Adoptive T Cell Therapy. Cancer Cell 2017;31(5):711–23 e4. doi: 10.1016/j.ccell.2017.04.003

48. Cao H, Dai P, Wang W, et al. Innate immune response of human plasmacytoid dendritic cells to poxvirus infection is subverted by vaccinia E3 via its Z-DNA/RNA binding domain. PLoS One 2012;7(5):e36823. doi: 10.1371/journal.pone.0036823 [published Online First: 2012/05/19]

49. Hildner K, Edelson BT, Purtha WE, et al. Batf3 deficiency reveals a critical role for CD8alpha+ dendritic cells in cytotoxic T cell immunity. Science 2008;322(5904):1097–100. doi: 10.1126/science.1164206 [published Online First: 2008/11/15]

50. Edelson BT, Kc W, Juang R, et al. Peripheral CD103+ dendritic cells form a unified subset developmentally related to CD8alpha+ conventional dendritic cells. J Exp Med 2010;207(4):823–36. doi: 10.1084/jem.20091627 [published Online First: 2010/03/31]

51. Zamarin D, Ricca JM, Sadekova S, et al. PD-L1 in tumor microenvironment mediates resistance to oncolytic immunotherapy. J Clin Invest 2018;128(4):1413–28. doi: 10.1172/JCI98047 [published Online First: 2018/03/06]

